# Optimisation of Weighted Ensembles of Genomic Prediction Models in Maize

**DOI:** 10.64898/2026.02.03.703660

**Authors:** Shunichiro Tomura, Owen Powell, Melanie J. Wilkinson, James Lefvre, Mark Cooper

## Abstract

Ensembles of multiple genomic prediction models have demonstrated improved prediction performance over the individual models contributing to the ensemble. The outperformance of ensemble models is expected from the Diversity Prediction Theorem, which states that for ensembles constructed with diverse prediction models, the ensemble prediction error becomes lower than the mean prediction error of the individual models. While a naïve ensemble-average model provides baseline performance improvement by aggregating all individual prediction models with equal weights, optimising weights for each individual model could further enhance ensemble prediction performance. The weights can be optimised based on their level of informativeness regarding prediction error and diversity. Here, we evaluated weighted ensemble-average models with three possible weight optimisation approaches (linear transformation, Nelder-Mead and Bayesian) using flowering time and tillering traits from two maize nested associated mapping (NAM) datasets; TeoNAM and MaizeNAM. The three proposed weighted ensemble-average approaches improved prediction performance in several of the prediction scenarios investigated. In particular, the weighted ensemble models enhanced prediction performance when the adjusted weights differed substantially from the equal weights used by the naïve ensemble models. For performance comparisons among the weighted ensembles, there was no clear superiority among the proposed approaches in both prediction accuracy and error across the prediction scenarios. Weight optimisation for ensembles warrants further investigation to explore the opportunities to improve their prediction performance; for example, integration of a weighted ensemble with a simultaneous hyperparameter tuning process may offer a promising direction for further research.

## Introduction

Plant breeders have a long history of research into new technologies for predictive modelling to accelerate genetic improvement of crops for agricultural systems (Cooper et al., 2009, 2021, 2025; Crossa et al., 2017, 2024). Genomic prediction (Meuwissen et al., 2001; Bernardo and Yu, 2007) has enhanced crop breeding programs by reducing breeding cycle length and consequently costs (Crossa et al., 2017; Voss-Fels et al., 2019). Instead of full dependency on phenotype records for selection, genomic prediction identifies potential candidates by predicting trait phenotypes from corresponding genomic marker values (Heffner et al., 2009; Escamilla et al., 2025). The performance of genomic prediction models is influenced by the complexity of the genetic architecture controlling target traits and the prediction algorithms, and hence various algorithms have been introduced to achieve higher prediction performance (Hammer et al., 2006; Riedelsheimer et al., 2012; Technow et al., 2015; Messina et al., 2018; Voss-Fels et al., 2019; Negus et al., 2024).

The success of each genomic prediction model has motivated the investigation of ensembles that aggregate the strengths of multiple genomic prediction models (Kick and Washburn, 2023; Cooper et al., 2025; Messina et al., 2025; Washburn et al., 2025). A range of ensemble-based prediction approaches have been introduced, employing both simple arithmetic averaging (Wallach et al., 2018; Kick and Washburn, 2023) and more complex aggregations (McCormick et al., 2021; Yan et al., 2021). The evaluation of these ensemble approaches revealed successful performance improvement across various traits and datasets. The potential of ensembles can be explained by the framework of the Many-Model Diversity Prediction Theorem (Hong and Page, 2004; Page, 2007, 2014, 2018), which states that the prediction error of an ensemble becomes lower than the mean error of individual prediction models, assuming that the ensemble consists of diverse prediction models. This theoretical framework provides a logical basis for the potential of ensemble approaches, explaining the improved genomic prediction performance in previous studies and thereby motivating the further investigation of ensemble algorithms.

In our previous investigations of ensembles (Tomura et al., 2025a, b, 2026), we demonstrated the framework of the Diversity Prediction Theorem as a foundation for constructing ensembles of prediction models that equally weigh the contributions of individual models. The observed results indicated inconsistent levels of performance improvement for different scenarios generated by various combinations of traits and datasets. The Diversity Prediction Theorem provides a possible interpretation of such inconsistency by emphasising the importance of model diversity in the construction of ensembles, when the individual models are given equal weight in their contribution to the ensemble. While the theorem does not directly inform the optimal weightings for models when their contributions to ensembles differ, it does provide a framework for exploring alternative weightings that can improve prediction performance relative to that of the naïve ensemble based on equal weights. This application of the Diversity Prediction Theorem is investigated as a novel contribution with broad applications for genomic prediction.

The weight adjustment problem of the ensembles shares a structural similarity with the framework of other optimisation problems. By adjusting key variables, optimisation approaches aim to achieve the best value of the performance criteria (highest benefit or lowest cost) in target complex problems. In the ensemble, a weight assigned to each individual prediction model can be a key variable, and the optimum weight values can be estimated based on performance criteria defined in the respective prediction algorithms. Through weight optimisation of the contributions of individual models, the predictive ability of ensembles was further improved for genomic prediction in animal breeding applications (Liang et al., 2021a, b; Yu et al., 2021; Liang et al., 2023; Wang et al., 2023). However, the potential of weight optimisation to improve performance of ensembles has not been well investigated in crop breeding applications.

Here, we investigated the potential to construct weighted ensemble-average models for genomic prediction in crop breeding using three different weight optimisation methods, two of which directly use the Diversity Prediction Theorem as a framework to identify weightings. The prediction performance was evaluated by extending our previous studies (Tomura et al., 2025a, b, 2026) focused on maize flowering time and tillering traits (Buckler et al., 2009; Chen et al., 2019); days to anthesis (DTA), anthesis and silking interval (ASI) and tiller number per plant (TILN). DTA and TILN are controlled by numerous genomic regions and their interactions within gene networks (Dong et al., 2012; Wisser et al., 2019; Powell et al., 2022; Hammer et al., 2023). Similarly, ASI is controlled by an extensive gene network, including the networks for the two flowering time traits (DTA and days to silking (DTS)) (Messina et al., 2019). Three objectives motivated this study: (1) Compare the prediction performance of the proposed weighted ensemble-average models against the benchmark ensemble using equal weights (naïve ensemble-average model) following the Diversity Prediction Theorem. (2) Analyse how the diversity in the individual genomic prediction models affected the prediction performance of the ensembles by applying the framework of the Diversity Prediction Theorem. (3) Compare the prediction performance of the three weighted ensemble-average models to identify which weighting approach consistently improved the prediction performance. Combining these three objectives, we apply the Diversity Prediction Theorem as a framework to investigate which weight optimisation method can outperform the naïve ensemble-average model by adjusting weights in proportion to the diversity of prediction performance at the individual prediction model level.

## Materials and Methods

### 1. Datasets

In this study, two maize nested association mapping (NAM) datasets were used to investigate the proposed ensemble approaches: the TeoNAM (Chen et al., 2019) and MaizeNAM (Buckler et al., 2009) datasets. The total number of recombinant inbred lines (RILs) and genomic markers (SNPs) was summarised in Table S1.

The TeoNAM dataset provides the genotype (genomic markers in the form of single nucleotide polymorphisms, SNPs) and trait phenotype records of RILs from five subpopulations, developed from hybrids between the maize inbred line W22 and five teosinte lines (*Z. mays ssp. parviglumis* for TIL01, TIL03, TIL11 and TIL14 and *Z. mays ssp. mexicana* for TIL25). Following the initial cross, the F1 was backcrossed once with W22. Subsequently, four generations were created through controlled self-pollination. The five RIL populations were evaluated twice through a randomised complete block design at the University of Wisconsin West Madison Agricultural Research Station. For W22TIL01, W22TIL03 and W22TIL11, the RILs were evaluated in the summers of 2015 and 2016. For W22TIL14, the RILs were evaluated in the summers of 2016 and 2017. W22TIL25 was evaluated at two different blocks in the summer of 2017. Since each subpopulation dataset consists of two record sets due to the evaluation of each RIL twice, both record sets were concatenated into one dataset by adding a factor column representing the respective environments in each subpopulation.

The MaizeNAM (Buckler et al., 2009) dataset provides genotype (SNPs) and trait phenotype records from RILs in 25 hybrid populations based on crosses between the maize inbred line B73 and 25 inbred lines generated from breeding programs operating in temperate and tropical regions. Following the initial cross to create the F1 generation, the F1 individual was self-pollinated to create the F2 generation. Each RIL population was generated by controlled self-pollination of a sample of F2 individuals until the generation was advanced to F5. The F5 RILs were tested twice in each of the four locations (Aurora in New York, Clayton in North Carolina, Columbia in Missouri and Urbana in Illinois) between 2006 and 2007 with a randomised design for each test (McMullen et al., 2009). The scored phenotypes for each RIL across the environments were used to calculate best linear unbiased predictors (BLUPs) with ASReml (v2.0) per population. The estimated BLUPs were linked to their corresponding genotypes as phenotype values.

The genetic diversity of the founders of the TeoNAM and MaizeNAM datasets was investigated by Hufford et al. (2012). The TeoNAM dataset is expected to have higher genetic diversity due to crosses with the ancestor species of maize (teosinte), introducing standing genetic variation at loci that have become fixed during the domestication process. In contrast, the MaizeNAM dataset is expected to have less genetic diversity due to crosses involving elite lines created post-domestication. The dataset diversity broadens the range of scenarios considered and is expected to affect the predictive performance of the individual genomic prediction models and the proposed weighted ensemble approaches. Hence, the difference in genetic marker pattern diversity between the TeoNAM and MaizeNAM datasets is advantageous for the objectives of this study.

Two flowering time-related traits (days to anthesis (DTA) and anthesis and silking interval (ASI)) from both datasets and the tiller number per plant (TILN) trait from the TeoNAM dataset were targeted for evaluation of the ensemble approaches. For DTA, the genetic architecture has been well-studied (Dong et al., 2012; Wisser et al., 2019), and hence it is possible to assess the model features for each ensemble approach at the genome level by comparing the estimated standing variation of the trait genetic architecture from each proposed approach with the well-known key gene regulators of DTA. The genetic architecture has also been actively investigated for TILN, revealing unique and complex biological interactions that are distinct from those of DTA (Powell et al., 2022; Hammer et al., 2023). ASI is a secondary trait that is measured as the difference between DTA and days to silking (DTS) flowering traits, potentially generating a more complex genetic architecture that is controlled by both traits and other developmental processes (Messina et al., 2019). Greater biological complexity of ASI might affect the prediction performance of the individual prediction models and proposed weighted ensemble approaches. The two maize NAM datasets and the three traits were used to investigate the predictive characteristics of the proposed weighted ensemble-average approaches.

### 2. Data preprocessing

Prior to the prediction performance evaluation of the ensembles, the two NAM datasets were processed by applying the three approaches leveraged in Tomura et al. (2025a). For the TeoNAM dataset, missing genetic markers were imputed using flanking markers. For the MaizeNAM dataset, Buckler et al. (2009) applied the flanking marker-based imputation, and hence no missing genomic markers were observed. RILs with missing phenotypes were removed from the TeoNAM dataset, while Buckler et al. (2009) had already removed RILs with missing phenotypes during the BLUP calculation process in the MaizeNAM dataset. Genomic markers were then filtered to mitigate the curse of dimensionality (Bellman, 1957; Ramstein et al., 2019), which can negatively affect the prediction patterns by containing an excessively large number of attributes (genomic markers, SNPs) compared to data points (RILs) for trait phenotype. Genomic markers with linkage disequilibrium (LD) higher than 0.8 were removed, applying a window size of 30,000 and a step size of 5 using PLINK (v1.9) (Chang et al., 2015) for the TeoNAM and MaizeNAM datasets.

### 3. Genomic prediction models

Three conventional genomic prediction models (ridge regression BLUP (rrBLUP), BayesB and reproducing kernel Hilbert Space regression (RKHS)) and three machine learning models (random forest (RF), support vector regression (SVR) and multi-layer perceptron (MLP)) were developed as individual genomic prediction models in this study using the computational tool EasiGP (Tomura et al., 2025b) in Python (v3.11.10) (Figure 1). rrBLUP and BayesB (Meuwissen et al., 2001) were implemented through a linear mixed model (Pérez and de Los Campos, 2014). These models are grounded in different assumptions regarding the distribution of genomic marker effects: normally distributed with the same variance for rrBLUP (Endelman, 2011), and a mix of zero effects and values from a t-distribution for BayesB. Such a difference in these assumptions is typically represented as shrinking a large amount of genomic marker effects to zero in BayesB, but not in rrBLUP. RKHS regression (Gianola and Van Kaam, 2008) uses a kernel based on squared Euclidean distance that implicitly maps data points (RILs) into a Hilbert space (de los Campos et al., 2009). RF (Breiman, 2001) is a tree-based approach consisting of a collection of decision trees that predict values by traversing a series of decision conditions from provided attributes (SNPs). The mean predicted values from each tree are calculated to determine the final prediction values. SVR (Drucker et al., 1996) optimises the location of a hyperplane by including the maximum number of data points within the space (epsilon tube) created between the hyperplane and support vectors. By optimising the hyperplane location based on data points mapped to higher dimensions using a kernel, SVR returns predicted values with nonlinear prediction patterns. MLP (Rosenblatt, 1962) predicts target values by mimicking the structure of the information transmission system in human brains. Numerous neurons are fully connected across several layers to aggregate information from others that were converted nonlinearly by applying an activation function. Consequently, the final prediction values are expected to be the sequential results of captured complex nonlinear prediction patterns.

**Figure 1:**
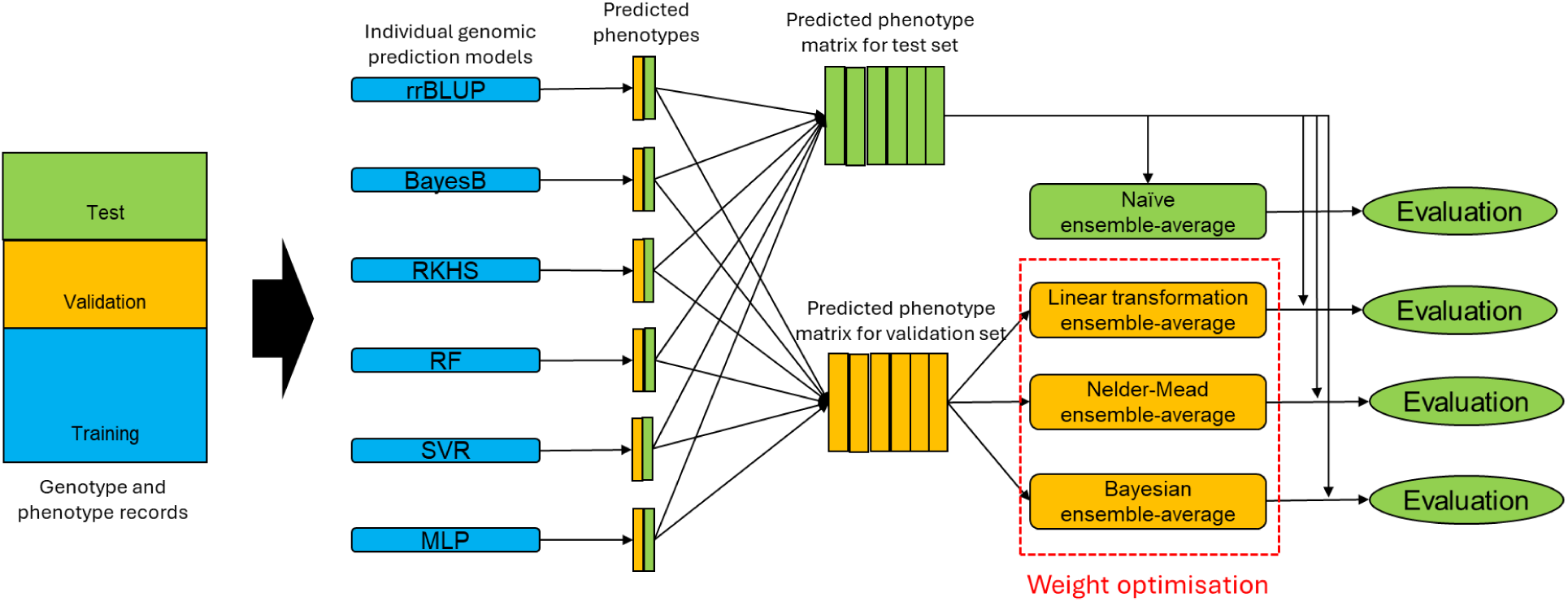
Model development and evaluation flow in this study. Genotype and phenotype records for each prediction scenario were split into training, validation and test sets, represented in blue, orange and green, respectively. The six individual genomic prediction models (rrBLUP, BayesB, RKHS, RF, SVR and MLP) were trained independently using the training set, followed by phenotype prediction on the validation and test sets. As a benchmark prediction performance, the naïve ensemble-average model was developed by calculating the mean predicted phenotypes for the test set using equal weights. The weights allocated to the individual genomic prediction models were optimised using the validation set through three optimisation approaches (linear transformation, Nelder-Mead and Bayesian ensemble-average). These weight optimisation approaches were expected to adjust weights assigned to each prediction model by maximising or minimising their objective function (Figure 3). The predicted phenotypes for the test set were weight-averaged using the identified weights. The three weighted ensemble-average models were evaluated against the naïve ensemble-average model (Figure 2, S2, S3).

For the conventional genomic prediction models (rrBLUP, BayesB and RKHS), the number of iterations was set to 12,000 and the burn-in was set to 2,000 in both datasets. The values for the other parameters were not adjusted from default settings. For rrBLUP, the Bayesian version of rrBLUP was developed and hence the number of iterations and burn-in were set. The number of decision trees was set to 1,000 in RF. Radial Basis Function (RBF) kernel was used in SVR. For the remaining hyperparameters in RF and SVR, the default hyperparameter values were leveraged for both datasets. In both datasets, MLP contained one hidden layer. For the TeoNAM dataset, 50 neurons were developed in the hidden layer with a dropout of 0, which were nonlinearly transformed using the Rectified Linear Unit (ReLU; Nair and Hinton, 2010) activation function. The constructed model was trained 200 times iteratively with a learning rate of 0.005 using the Adaptive Moment Estimation with decoupled weight decay (AdamW; Loshchilov and Hutter, 2017) optimiser. For the MaizeNAM dataset, the number of neurons was set to 10 with a dropout of 0.1. ReLU was also used as the activation function. The prediction model was trained with 2,500 iterations with a learning rate of 0.005 using the Root Mean Square Propagation (RMSprop; Tieleman, 2012) optimiser. Different hyperparameter settings were used for each dataset since the selected hyperparameter setting for one dataset achieved considerably low prediction performance on the other. These hyperparameter settings for each prediction model were previously demonstrated to provide stable results for applications of these prediction methods in the TeoNAM and MaizeNAM datasets (Tomura et al., 2025a, b, 2026). The combinations of hyperparameter values were selected heuristically from a range of options, and hence other hyperparameter value combinations might possibly achieve higher prediction performance. Hyperparameter tuning in consideration with weight optimisation methods can be a research extension for weighted ensemble studies, as mentioned later in the Discussion.

### 4. Weight optimisation

Three weight optimisation approaches (linear transformation, Nelder-Mead and Bayesian) were implemented to construct weighted ensembles as a part of the EasiGP software pipeline (Tomura et al., 2025b) operation in this study (Figure 1).

The linear transformation ensemble-average model was implemented to discover the set of optimum weights that minimises the mean squared error (MSE) objective function using the neural network approach (Figure S1a). A set of predicted phenotypes from each individual genomic prediction model was multiplied by a corresponding trainable weight. The weighted predicted trait phenotypes from all individual genomic prediction models were summed as the final predicted phenotypes. The optimum value for the weights was determined through the iterative training process. The weight values were adjusted in each iteration to minimise the difference between the predicted and observed trait phenotype values. However, the randomness in the initial weight values affected the ideal number of training iterations, and hence the standard training approach could not effectively identify the optimum weights. To mitigate this problem, the linear transformation ensemble-average model was equipped with an early stop. If the prediction error (MSE) for the validation set did not improve after *M* iterations (defined as patience), the training process was terminated. After the development of the *N* independent ensemble models with the early stop, the one that reached the lowest prediction error in the validation set was selected as the final linear transformation ensemble-average model. The total number of neurons was set to six (the total number of individual genomic prediction models in this study was six). In both datasets, the total number of epochs was 150 with the optimiser of Adaptive Moment Estimation (Adam; Kingma and Ba, 2014). The learning rate was 0.005 with the weight decay of 0.01. The size of each batch was set to 2. The number of the independent linear transformation ensemble-average model (*N*) and patience (*M*) were set to 30 and 10, respectively.

The Nelder-Mead ensemble-average model heuristically determined the optimum set of weights that minimised an objective function using the Nelder-Mead algorithm (Nelder and Mead, 1965) (Figure S1b). The algorithm randomly selected sets of values for the vertices (weights in this study) prior to the implementation. Those sets were defined as Simplex, and an output value from the objective function using each vertex was returned. Subsequently, the vertices were ranked by the value of the returned output, and the one returning the worst performance in relation to the objective function was replaced with another vertex calculated based on the concept of reflection, expansion and contraction (Singer and Nelder, 2009). This replacement process was repeated until the output values from the objective function converged or until the defined number of iterations was reached. The objective function used in this study was derived from the Diversity Prediction Theorem (Page, 2018) defined below:

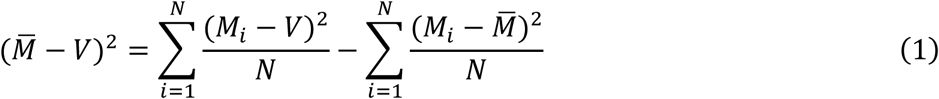

where *M_i_* is the predicted value from the prediction model *i*, M̅ is the mean predicted value from the *i* individual prediction models calculated from the arithmetic mean operation with the same weight, *V* is the true value and *N* is the total number of prediction models included in the ensemble mechanism. The Many-Model error (first term) is estimated by the subtraction of the prediction diversity (third term) from the mean error of individual prediction models (second term). In this study, observed phenotypes were defined as the true value *V*, and the Many-Model error was leveraged as the ensemble error. The Diversity Prediction Theorem indicates that the diversity in the predicted values can reduce the ensemble prediction error compared to the mean prediction error of individual prediction models. The minimisation of Equation (1) can improve the prediction performance of the ensemble, achieved by adjusting the contribution of each individual genomic prediction model to each term value using weights. Hence, the objective function for the Nelder-Mead ensemble-average model can be represented below:

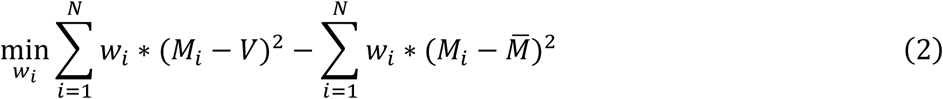

where *w_i_* is the weight allocated to the individual prediction model *i*. When the weights were equal as applied in the naïve ensemble-average model, the weights were represented as constant: 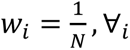. In the weighted ensembles, the weights were adjusted to minimise Equation (2). The weight values were optimised to increase the prediction diversity values for minimising the ensemble error in relation to the mean error of the individual prediction models. In both datasets, the total number of weights was six (the total number of individual genomic prediction models). Weight values were sought within the range of 0 and 10 and then normalised. The threshold for the convergence was 1*10^-8^. The other parameters were set to default.

The Bayesian ensemble-average model statistically estimated the optimum variable (weight) values to maximise an objective function (Figure S1c). Using the values of the objective function based on the random sets of weight values, the algorithm estimated the ranges of potential sets of weight values that are likely to be the optimum by developing a surrogate model. An acquisition function converted the information from the surrogate model to estimate the next set of weights that was likely to maximise the objective function. This process of sampling and updating was iterated until the returned value converged or reached the end of the iteration process. The objective function for the Bayesian ensemble-average model was defined below based on Equation (2):

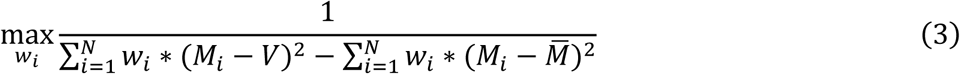

The weights (*w*) were adjusted to maximise Equation (3). Since Equation (3) is the inverse of Equation (2), the maximisation of Equation (3) leads to the minimisation of Equation (2). In both datasets, the total number of weights was six, which were optimised within the range of values 0 and 10 with normalisation. The expected improvement was used as the acquisition function with the exploration-exploitation trade-off hyperparameter of 1*10^-6^. The other hyperparameters were set to default.

The prediction performance of these three weighted ensemble-average models was compared with the naïve ensemble-average as a benchmark. Following the Diversity Prediction Theorem, the naïve ensemble-average model (Tomura et al., 2025a) calculated the arithmetic mean of predicted trait phenotypes from all individual genomic prediction models with equal weights.

### 5. Genomic marker and marker-by-marker interaction effect inference

Genomic marker (SNP) effects were extracted from the individual genomic prediction models and the ensemble models using the back-calculation functions in EasiGP (Tomura et al. 2025b). For rrBLUP and BayesB, the allele substitution effects were used as the SNP effects. For RKHS, SVR, MLP and pairwise SNP effects (genomic marker-by-marker interaction effects) from RF, Shapley scores (Shapley, 1953; Lundberg and Lee, 2017) were used to compare the causal effect of a specific SNP represented in an accumulative conditional effect. Shapley scores from the 50 randomly chosen data points (RILs) were leveraged as the SNP effects in the absolute value format for each prediction scenario. For RF, the impurity-based approach (Ishwaran, 2015) was applied to extract SNP effects. A higher importance (SNP effect size) value was allocated to the corresponding attribute (SNP) if the SNP value clearly clusters data points (RILs) with similar target (trait phenotype) values. The SNP effects for the naïve ensemble-average model were estimated by calculating the mean value of the normalised SNP effects from the individual genomic prediction models. For the weighted ensemble models, the mean value of the normalised SNP effects from all the individual genomic prediction models was calculated with weights estimated in each method.

### 6. Visualisation of the standing variation of the trait genetic architecture

The extracted genomic marker SNP effects and interactions from the genomic prediction models were mapped to respective genomic marker regions using circos plots (Krzywinski et al., 2009) to visualise the standing variation of the trait genetic architecture derived from each individual prediction model and the four ensemble-based models as the final phase in EasiGP operations (Tomura et al., 2025b). The first few rings of the circos plot represented key genome regions identified in the previous studies. For the circos plots of DTA, the innermost ring indicated QTL regions from Chen et al. (2019) for the TeoNAM dataset and Buckler et al. (2009) for the MaizeNAM dataset. The second innermost ring showed the location of each QTL from Wisser et al. (2019). The third and fourth rings represented the key gene locations that operate on the leaf and shoot apical meristem (SAM), respectively, from Dong et al. (2012). The subsequent rings represented the estimated SNP effects from each genomic prediction model. For ASI, QTL information (Chen et al. (2019) for the TeoNAM dataset and Buckler et al. (2009) for the MaizeNAM dataset) was used as the key gene region ring, and the SNP genomic marker effect rings followed afterwards. For the TeoNAM and MaizeNAM datasets, the range of each SNP genomic marker on the circos plot was extended by 0.2 cM and 1 cM, respectively, on each side for visualisation purposes. Internal links between SNP positions illustrated genomic marker pairs with the top 0.01% of the highest pairwise Shapley scores in both datasets.

### 7. Genomic prediction model evaluation

Two metrics were used to evaluate the prediction performance of each genomic prediction model. Pearson correlation measured the prediction accuracy, whereas MSE measured the prediction error. For the calculation of the diversity level in each proposed ensemble approach, the ratio of the prediction diversity (third term) over the mean error of individual prediction models (second term) from the Diversity Prediction Theorem was leveraged by updating Equation (1):

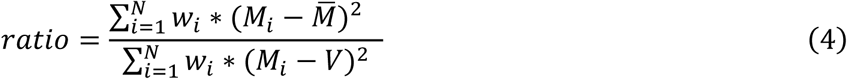

Since the scale of each phenotype differed from traits and datasets, the direct comparison of the third term in Equation (1) did not allow for a balanced comparison of the diversity observed in each ensemble approach. Additionally, the level of diversity needs to be analysed in relation to the mean error of individual genomic prediction models. The Many-Model error (first term) would still be high even if the prediction diversity (third term) was high when the mean error of individual prediction models (second term) was also high. A higher ratio value in Equation (4) indicated more diversified phenotypes in relation to the second term in Equation (1).

The prediction performance and the diversity level of the four ensemble models were iteratively evaluated under various prediction scenarios (Figure 1). For the TeoNAM dataset, each subpopulation data was sampled 500 times, and hence 2,500 (5 populations * 500 samples) different prediction scenarios were generated per trait. For the MaizeNAM dataset, each subpopulation data was sampled 50 times, and hence 1,250 (25 populations * 50 samples) different prediction scenarios were generated per trait. The data were split into training, validation and test sets with the ratio of 0.5-0.25-0.25 (Figure 1). The optimum weights were estimated from the validation set. The prediction performance of each ensemble approach was evaluated using the test set in each prediction scenario. The SNP genomic marker and marker-by-marker interaction effects for the circos plots were estimated by calculating the mean values across all the prediction scenarios in each trait.

## Results

### 1. The weighted ensemble-average models improved prediction performance for DTA and TILN

The prediction ability of the proposed weighted ensemble models depended on the target traits. For DTA, all weighted ensemble-average models reached higher median prediction accuracy and lower median prediction error than the naïve ensemble model, especially for the TeoNAM dataset (Figure 2, S2, S3; Table S2). In the TeoNAM dataset, the Nelder-Mead ensemble model reached the highest median prediction accuracy (Pearson correlation = 0.879) and the lowest median prediction error (MSE = 8.448). Similarly, the Nelder-Mead ensemble model achieved the highest median prediction accuracy (0.625) and lowest median prediction error (2.436) in the MaizeNAM dataset, with a small performance difference compared to other ensemble-average models. Such superiority of the weighted ensemble models was also observed when compared with the individual genomic prediction models (Figure S2; Table S2). These results indicate that the weightedensemble-average models can improve prediction performance for DTA in both the TeoNAM and MaizeNAM datasets.

**Figure 2:**
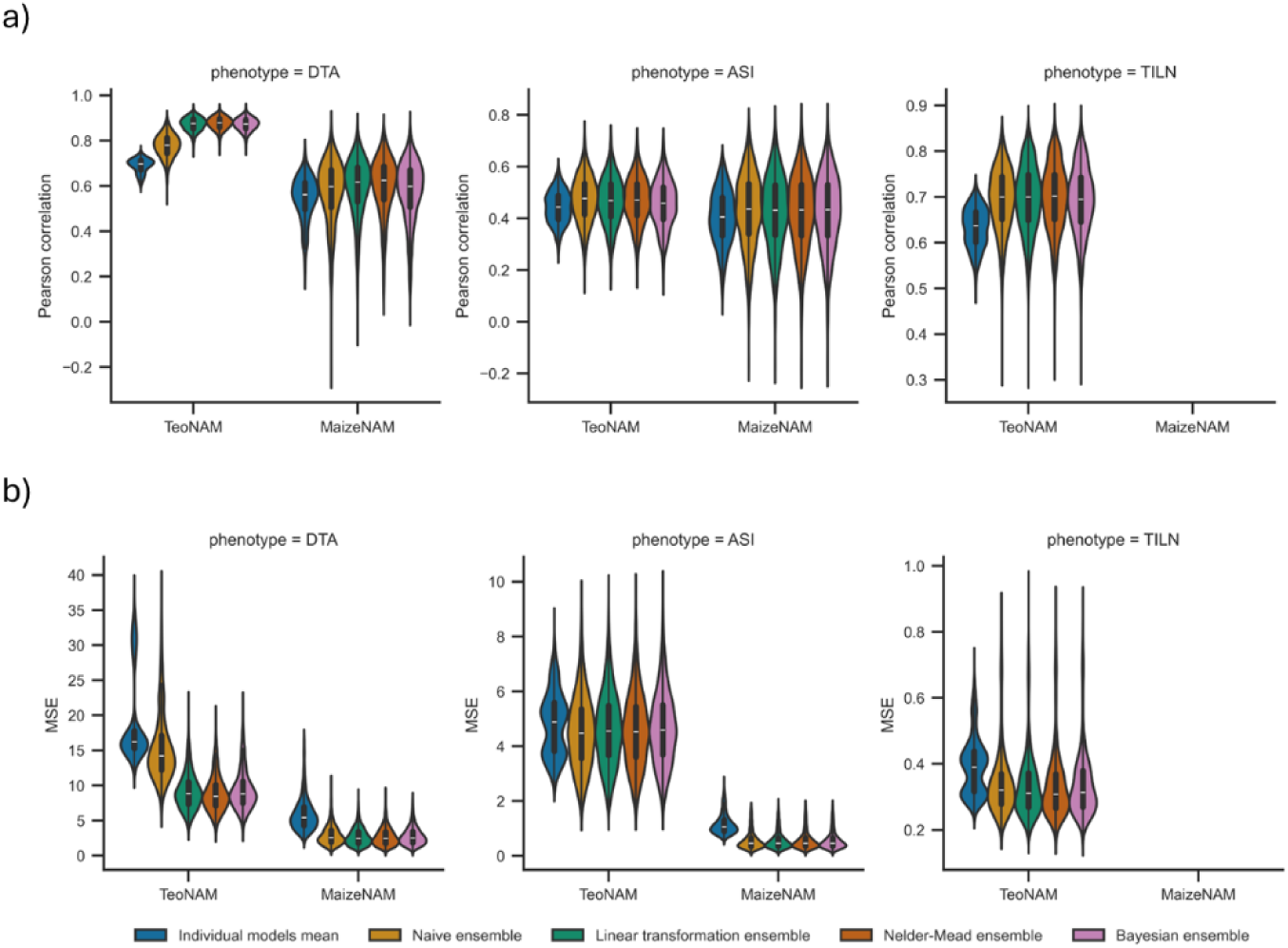
Comparison of median prediction performance between the mean metric value of the individual genomic prediction models (rrBLUP, BayesB, RKHS, RF, SVR and MLP) and the ensemble-average models (naïve ensemble, linear transformation ensemble, Nelder-Mead ensemble and Bayesian ensemble) for the days to anthesis (DTA), anthesis and silking interval (ASI) and tiller number per plant (TILN) traits in the TeoNAM and MaizeNAM datasets. The TILN trait was recorded only in the TeoNAM dataset. The prediction performance was measured using the two metrics: a) Pearson correlation and b) mean squared error (MSE). The prediction metrics were calculated from 2,500 prediction scenarios for the TeoNAM dataset and 1,250 prediction scenarios for the MaizeNAM dataset for each trait. The width of the violins represents the distribution of performance metrics. The white horizontal lines on the black box plots show the median value for each metric. The whiskers extend 1.5 times the interquartile range. The median value of each metric is recorded in Table S2.

For TILN, the prediction performance improvement using the weighted ensemble-average models was mainly observed in the prediction error (Figure 2, S2, S3; Table S2). While the Nelder-Mead ensemble also achieved the highest median prediction accuracy (0.702), no clear performance improvement over the naïve ensemble-average model (0.700) was observed. In contrast, the Nelder-Mead ensemble reached the lowest median prediction error (0.308), showing a noticeable difference from the median prediction error of the naïve ensemble-average model (0.321).

For ASI, however, such outperformance of the weighted ensemble-average models was not clearly observed. The naïve and weighted ensemble methods performed similarly (Figure 2, S2, S3; Table S2). In the TeoNAM dataset, while the naïve ensemble-average model reached the highest median prediction accuracy (0.477) and lowest median prediction error (4.476), the difference from the other ensemble models was subtle (prediction accuracy ranging from 0.459 to 0.471; prediction error ranging from 4.523 to 4.589). In the MaizeNAM dataset, the prediction performance difference between the naïve ensemble-average model and the weighted ensemble-average models was negligible in both median prediction accuracy (ranging from 0.432 to 0.436) and prediction error (ranging from 4.461 to 4.666). Only small differences in the prediction performance were observed when the ensembles were compared with the individual genomic prediction models (Figure S2; Table S2). The weighted ensemble-average model did not clearly improve the prediction performance of the naive ensemble average model for ASI in either the TeoNAM or MaizeNAM dataset.

### 2. Individual models contributed differently to the weighted ensembles across the traits and datasets

The contribution level of each individual genomic prediction model for the proposed weighted ensemble models highly depended on the traits. For DTA, the parametric and semiparametric genomic prediction models (rrBLUP, BayesB and RKHS) were assigned heavier weights with more diverse weighting patterns compared to the machine learning models (RF, SVR and MLP) (Figure 3a; Table S3a). For the TeoNAM dataset, the mean sum of weights from the parametric and semiparametric genomic prediction models across the weighted ensembles (0.77) was considerably higher than that from the machine learning models (0.23). The differences between the highest and lowest normalised mean weights for the parametric and semiparametric genomic prediction models were also higher than the machine learning models (rrBLUP = 0.17, BayesB = 0.32, RKHS = 0.29, RF = 0.10, SVR = 0.01, MLP = 0.05). The same trend was observed in the MaizeNAM dataset, showing heavier mean summed weights from the parametric and semiparametric genomic prediction models across the three weighted ensembles (0.69) than the machine learning models (0.31), with higher weight value differences (rrBLUP = 0.21, BayesB = 0.08, RKHS = 0.34, RF = 0.13, SVR = 0.09, MLP = 0.08). This comparison indicates that the proposed weighted-ensemble approaches emphasised predictive information from the parametric and semiparametric genomic prediction models more with diverse weighting patterns for DTA.

**Figure 3:**
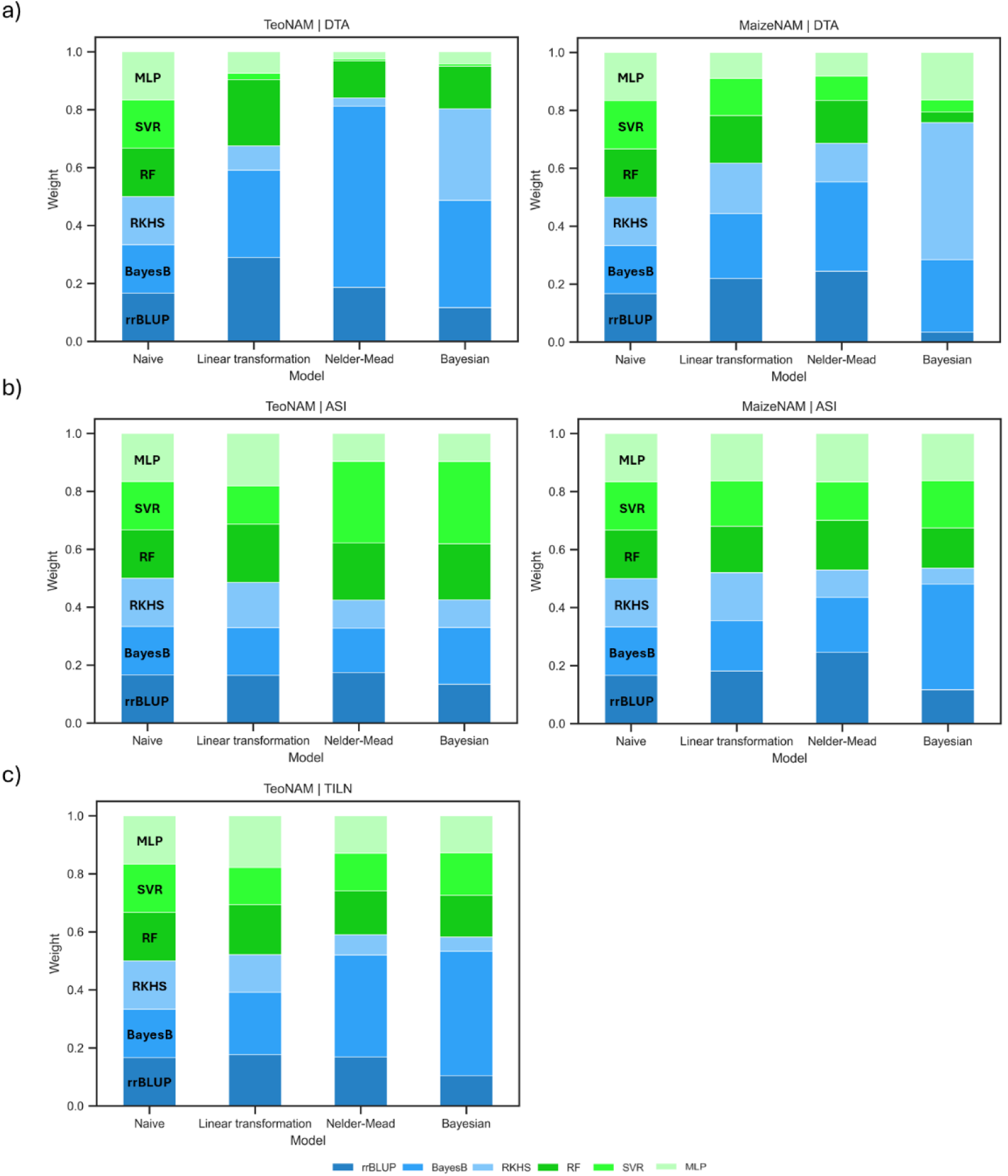
The comparison of mean weights allocated to the six individual genomic prediction models (rrBLUP, BayesB, RKHS, RF, SVR and MLP) by the naïve ensemble-average model and the three weighted ensemble-average models (linear transformation, Nelder-Mead and Bayesian) for a) the days to anthesis (DTA), b) anthesis and silking interval (ASI) and c) tiller number per plant (TILN) traits in the TeoNAM and MaizeNAM datasets. The TILN trait was recorded only in the TeoNAM dataset. The blue colours represent weights for parametric and semiparametric models (rrBLUP, BayesB and RKHS), while the green colours indicate weights for machine learning models (RF, SVR and MLP). The weights were extracted from 2,500 prediction scenarios for the TeoNAM dataset and 1,250 prediction scenarios for the MaizeNAM dataset. The actual mean weight and standard error values are reported in Table S3.

In contrast, heavier weights were assigned to the machine learning models with similar weighting patterns across the three weighted ensemble-average models for ASI (Figure 3b; Table S3b). The mean summed weight of parametric and semiparametric genomic prediction models across the weighted ensemble models (0.45) became smaller than that of machine learning models (0.56) for the TeoNAM dataset. The weight value differences in the parametric and semiparametric genomic prediction models became smaller for ASI compared to DTA (rrBLUP = 0.04, BayesB = 0.04, RKHS = 0.06, RF = 0.01, SVR = 0.15, MLP = 0.09). The same trend was observed for the MaizeNAM dataset, with increased mean summed weight from the machine learning models (0.47) that was comparable to that from the parametric and semiparametric models (0.53), with less diverse weighting patterns than DTA (rrBLUP = 0.13, BayesB = 0.19, RKHS = 0.11, RF = 0.03, SVR = 0.03, MLP = 0.00). These observed results indicate that the predictive information from the machine learning models was emphasised more for ASI, and the naïve ensemble-average model weights were within the range of values identified by all three weighted ensemble algorithms. A smaller deviation in the weight combination identified by the weighted ensemble-average models from that of the naïve ensemble-average model led to similar prediction performance in both naïve and weighted ensemble-average models for ASI (Figure 2, 3, S2, S3; Table S2, S3).

For TILN, heavier weights on the parametric and semiparametric models were also observed but less emphasised compared to DTA (Figure 3c; Table S3c). For the TeoNAM dataset, the mean sum of weights from the parametric and semiparametric genomic prediction models across the weighted ensembles (0.57) was higher than that from the machine learning models (0.43). However, the difference between the two groups was reduced when compared with DTA. The differences between the highest and lowest normalised mean weights for the parametric and semiparametric genomic prediction models were higher than the machine learning models (parametric and semiparametric models ranging from 0.08 to 0.21, machine learning models ranging from 0.02 to 0.05). Overall, TILN exhibited intermediate values between DTA (heavier and diverse weights on the parametric and semiparametric models) and ASI (more equal and similar weights on all individual prediction models).

### 3. The Diversity Prediction Theorem provided insight into when weight optimisation reduces prediction error

The framework of the Diversity Prediction Theorem revealed that the effect of the weight optimisation on diversity at the individual level was highly trait-dependent. The Many-Model error (first term) for the weighted ensemble-average models was lower than that of the naïve ensemble-average models for DTA and TILN across the datasets (Table 1). This lower Many-Model error was attributed to reduced mean prediction error (second term) and enhanced model diversity (third term) at the individual level compared to those of the naïve ensemble-average models. Consequently, the ratio of the third term over the second term tended to be considerably higher for DTA and TILN. The Bayesian ensemble-average model achieved the highest ratio value for these traits in both datasets. In contrast, no notable differences in the values of each term were observed for the ASI of both datasets. While the naïve ensemble-average and the Bayesian ensemble-average models achieved the highest ratio values for the TeoNAM and MaizeNAM datasets, respectively, the differences among the ensemble models were subtle. These results indicated that when the three weighted ensemble-average models did not diversify the individual genomic prediction models more than the naïve ensemble-average model, they failed to improve prediction performance relative to the naïve ensemble-average model.

**Table 1:** Association analysis between the diversity level of individual prediction models and prediction error of ensembles. The mean value of each term in the Diversity Prediction Theorem from the three proposed weighted ensemble-average models (linear transformation, Nelder-Mead and Bayesian) was compared with that of the naïve ensemble-average model. The first, second and third represent the Many-Model (ensemble) error, Average-Model error and diversity of model predictions, respectively. The third/second term is the ratio of the mean prediction diversity over the mean average error of the individual prediction models. For the first and second terms, the lowest value was highlighted in bold in each trait-by-dataset prediction scenario. For the third and third/second term, the highest value was highlighted in bold in each trait-by-dataset prediction scenario. The mean value for each term was calculated for the days to anthesis (DTA), anthesis and silking interval (ASI) and tiller number per plant (TILN) traits in the TeoNAM and MaizeNAM datasets. The TILN trait was recorded only in the TeoNAM dataset.

| Trait | Term | TeoNAM |  |  |  | MaizeNAM |  |  |  |
| --- | --- | --- | --- | --- | --- | --- | --- | --- | --- |
|  |  | Naïve | Linear transformation | Nelder-Mead | Bayesian | Naïve | Linear transformation | Nelder-Mead | Bayesian |
| DTA | Many-Model error (first term) | 22.090±0.044 | 10.515±0.037 | 4.822±0.032 | **3.631±0.026** | 3.618±0.019 | **2.703±0.018** | 2.831±0.018 | 3.143±0.018 |
|  | Average-Model error (second term) | 33.544±0.053 | 24.391±0.047 | 18.424±0.038 | **16.351±0.028** | 5.975±0.023 | **4.269±0.022** | 4.391±0.021 | 5.322±0.023 |
|  | Diversity of model predictions (third term) | 11.455±0.014 | **13.876±0.022** | 13.602±0.021 | 12.720±0.019 | **2.357±0.010** | 1.566±0.008 | 1.560±0.007 | 2.179±0.010 |
|  | third term / second term | 0.341 | 0.569 | 0.738 | **0.778** | 0.394 | 0.367 | 0.355 | **0.409** |
| ASI | Many-Model error (first term) | 5.227±0.015 | **5.213±0.015** | 5.255±0.015 | 5.292±0.016 | 0.597±0.003 | 0.582±0.003 | 0.583±0.003 | **0.576±0.003** |
|  | Average-Model error (second term) | 6.122±0.015 | **6.033±0.015** | 6.058±0.015 | 6.088±0.016 | 0.667±0.003 | 0.653±0.003 | 0.655±0.003 | **0.649±0.003** |
|  | Diversity of model predictions (third term) | **0.895±0.002** | 0.819±0.002 | 0.803±0.002 | 0.796±0.002 | 0.071±0.000 | 0.071±0.000 | 0.072±0.000 | **0.073±0.000** |
|  | third term / second term | **0.146** | 0.136 | 0.133 | 0.131 | 0.106 | 0.109 | 0.109 | **0.112** |
| TILN | Many-Model error (first term) | 0.482±0.001 | 0.447±0.001 | 0.414±0.001 | **0.311±0.001** | - | - | - | - |
|  | Average-Model error (second term) | 0.633±0.001 | 0.606±0.001 | 0.571±0.001 | **0.471±0.001** | - | - | - | - |
|  | Diversity of model predictions (third term) | 0.151±0.000 | 0.159±0.000 | 0.157±0.000 | **0.160±0.000** | - | - | - | - |
|  | third term / second term | 0.239 | 0.262 | 0.275 | **0.340** | - | - | - | - |

### 4. The ensemble models returned similar predicted phenotypes and the standing variation of the trait genetic architecture

The differences between naïve ensemble-average and the three weighted ensemble-average models were small at both predicted phenotype and genome levels. The pairwise comparison of the ensemble models at the predicted phenotype level showed a strong mean Pearson correlation across pairs, especially in the MaizeNAM dataset (DTA = 0.990, ASI = 0.982) compared to the TeoNAM (DTA = 0.939, ASI = 0.931, TILN = 0.989) dataset for both traits (Figure 4, S4). The stronger correlation for the MaizeNAM dataset was highlighted when the mean correlation was calculated for the pairs with the naïve ensemble-average model, especially for DTA (MaizeNAM = 0.985, TeoNAM = 0.901) rather than ASI (MaizeNAM = 0.973, TeoNAM = 0.953). While the observed correlations between the naïve ensemble-average model and the weighted ensemble-average models were high for both datasets and traits, the lower correlation with the naïve ensemble-average model for DTA in the TeoNAM dataset corresponded to the larger improvement of prediction performance using the weighted ensemble-average models for DTA in the TeoNAM dataset.

**Figure 4:**
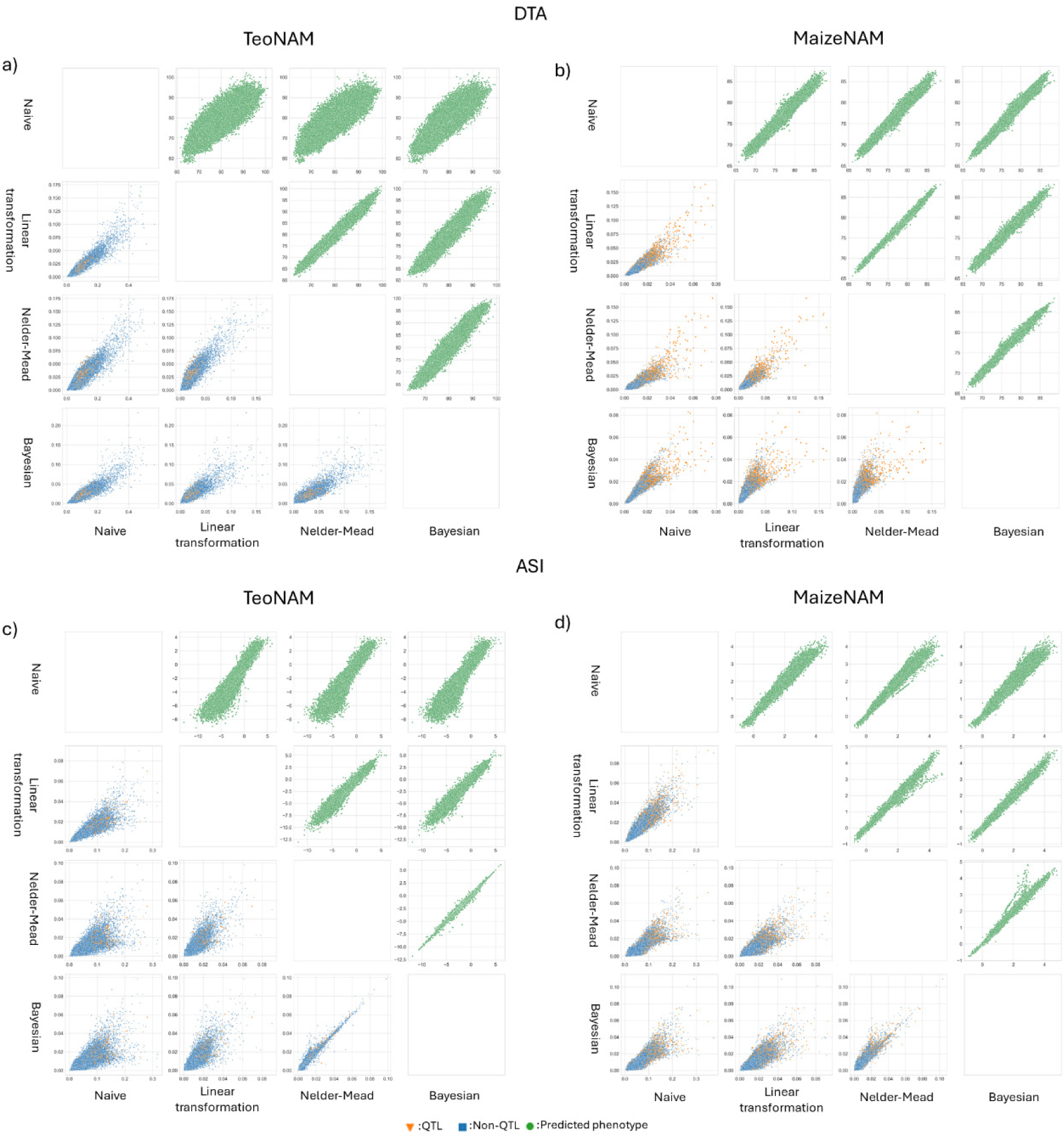
Pairwise comparisons of the ensemble models (naïve ensemble, linear transformation, Nelder-Mead and Bayesian) for the days to anthesis (DTA) and anthesis and silking interval (ASI) trait across all the prediction scenarios for the TeoNAM (2,500 prediction scenarios) and MaizeNAM (1,250 prediction scenarios) datasets; a) DTA for the TeoNAM dataset, b) DTA for the MaizeNAM dataset, c) ASI for the TeoNAM dataset and d) ASI for the MaizeNAM dataset. Within each scatter plot matrix, the ensemble models were compared at the predicted phenotypes (top right triangle) and genomic marker effects (the bottom left triangle) levels. The green dots represent a pair of predicted phenotypes for RIL samples in the test sets for each prediction scenario. The blue squares and orange triangles represent pairs of inferred effects of genomic markers in each sample scenario for non-QTL and QTL, respectively. Non-QTL and QTL markers were identified by Chen et al. (2019) for the TeoNAM dataset and Buckler et al. (2009) for the MaizeNAM dataset.

Similarly, the mean correlation values for the genomic marker effects in the MaizeNAM (DTA = 0.926, ASI = 0.921) and TeoNAM (DTA = 0.929, ASI = 0.931, TILN = 0.931) datasets were high. This strong correlation between the ensemble pairs was observed as clear positive associations for genomic markers with small effect sizes, while weaker correlations between the pairs were observed for genomic markers assigned larger effect sizes (Figure 4, S4). These association patterns indicate that more variation in quantification patterns for genomic markers with large effect sizes resulted in slightly more contrasting predicted phenotypes, especially in the TeoNAM dataset. Nonetheless, both predicted phenotypes and genomic marker effects of the ensemble approaches were similar, considering the high mean correlation values.

The similarity of the weighted ensemble-average models at the genome level was also observed in the constructed circos plots representing the inferred standing variation of the genetic architecture of DTA, ASI and TILN (Figure 5, S5). Each weighted ensemble-average model highlighted similar genome regions as highly influential to the target traits across the chromosomes. These consistent quantification patterns were emphasised when compared with the inferred standing variation of the trait genetic architecture from the individual genomic prediction models that showed highly diverse patterns of genomic marker effects, especially for the TeoNAM dataset (Figure 5, S5). This indicates that the proposed ensemble models captured similar standing variation of the trait genetic architectures across the chromosomes for the target traits, especially in the MaizeNAM dataset.

**Figure 5:**
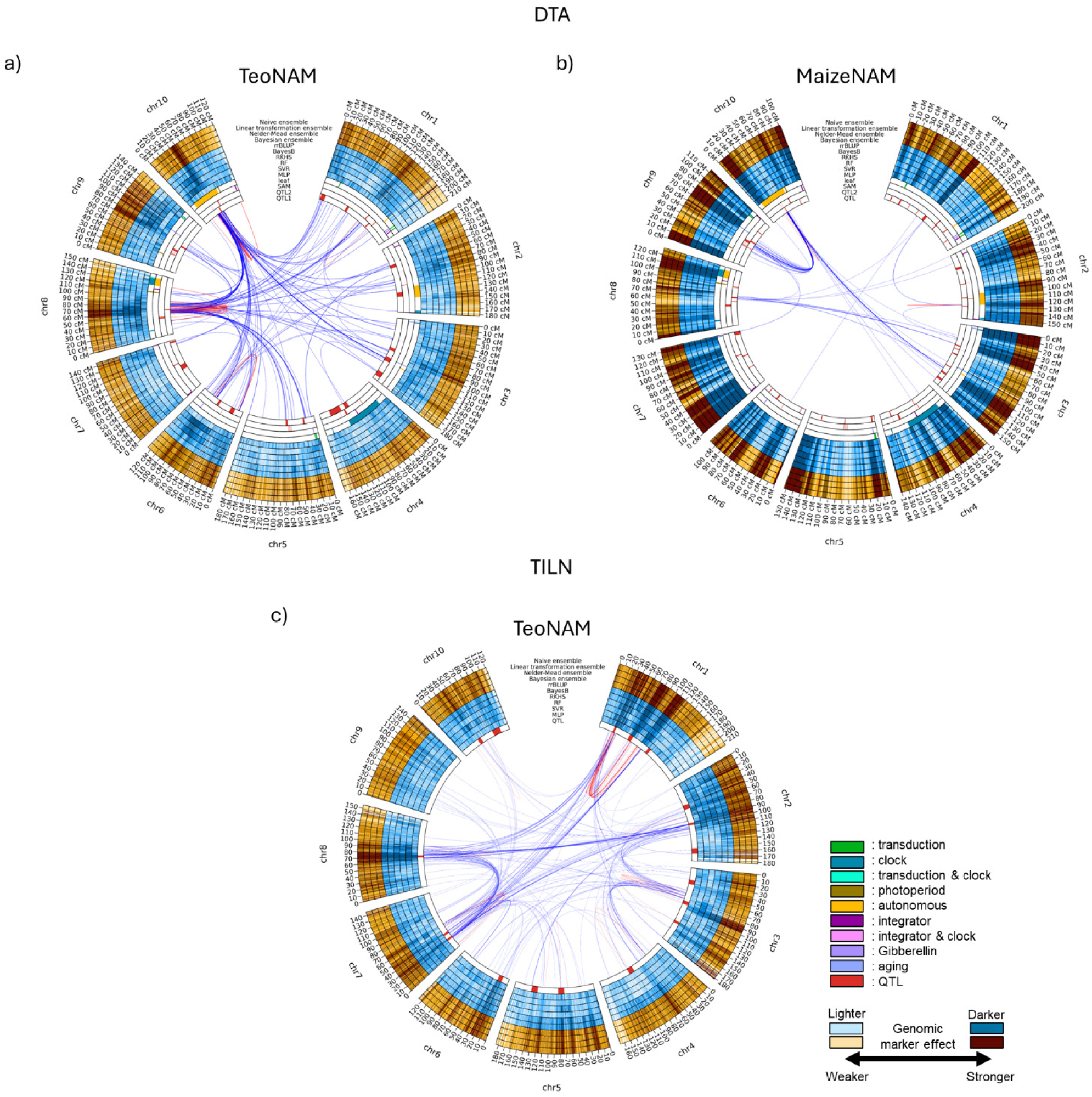
Circos plots for the days to anthesis (DTA) and tiller number per plant (TILN) trait in the TeoNAM and MaizeNAM datasets; a) DTA for the TeoNAM dataset, b) DTA for the MaizeNAM dataset and c) TILN for the TeoNAM dataset. The TILN trait was recorded only in the TeoNAM dataset. For DTA, the innermost (QTL 1) ring represents the QTL gene regions estimated by Chen et al. (2019) for the TeoNAM dataset and by Buckler et al. (2009) for the MaizeNAM dataset, respectively. The second innermost ring (QTL 2) represents the QTL gene regions identified by Wisser et al. (2019). The third and fourth innermost rings indicate the locations of gene regulators for maize flowering time that affect the shoot apical meristem (SAM) and leaf, respectively, based on Dong et al. (2012). For TILN, the innermost (QTL) ring shows the QTL gene regions estimated by Chen et al. (2019) for the TeoNAM dataset. The blue rings indicate genomic marker effects across the gene regions estimated by MLP, SVR, RF, RKHS, BayesB and rrBLUP. The subsequent four orange rings outwards are the genomic marker effects estimated by Bayesian, Nelder-Mead, linear transformation and naïve ensemble. The darkness level of the blue and orange colours indicates the strength of the genomic marker effects, sectioned into ten levels using the quantiles. Darker colours represent higher genomic marker effect sizes. The red and blue lines between genome regions show the genomic marker interaction effects calculated from pairwise Shapley scores from RF (top 0.01%; red = within chromosome and blue = between chromosomes).

### 5. The weighted ensemble models highlighted genome regions containing key genes for the target traits

Several highlighted genomic marker regions contributing to the standing variation associated with the trait genetic architecture for each ensemble model overlapped with key gene regulators identified in previous studies (Figure 5, S5). For DTA in the TeoNAM dataset (Figure 5a), the genome region between 35 and 50 cM in chromosome 10 was repeatedly highlighted by the ensemble models. This region overlaps with *ZmCCT10*, upregulating the circadian clock by controlling the photoperiod pathway (Dong et al., 2012; Chen et al., 2019; Wisser et al., 2019). Another example was the region between 60 and 70 cM, which was consistently highlighted by the ensemble models, with heavy interactions with *ZmCCT10* identified by RF. This region contains *ZCN8*, which regulates maize flowering time through interactions with *ZmCCT10;* another key gene for the photoperiod pathway (Dong et al., 2012; Chen et al., 2019; Wisser et al., 2019). In the MaizeNAM dataset (Figure 5b), the genome region between 50 and 60 cM was highlighted by the ensemble models in addition to *ZmCCT10* and *ZCN8*. This region overlapped with the QTL region containing *ZmCCT9*, which is another photoperiod gene that controls *ZCN8* with a negative effect (Huang et al., 2018; Wisser et al., 2019). For TILN (Figure 5c), the ensemble models highlighted the narrow genome region at 125 cM in chromosome 1. This region was close to the genome region containing *TB1*, which is one of the most significant genes involved in tillering of maize. *TB1* regulates the growth of axillary meristems that become tillers (Doebley et al., 1997; Chen et al., 2019). Additionally, the ensemble models featured the genome region between 85 and 95 cM in chromosome 3. This region contains *Zea AGAMOUS2* (*ZAG2*) that regulates tillering downstream of *TB1* as a MADS-box gene (Studer et al., 2017; Chen et al., 2019). Another member of the MADS-box gene family was also highlighted in chromosome 1 (*Zmm20*; between 30 and 40 cM) (Zhao et al., 2011; Chen et al., 2019). The ensemble models also emphasised *PROSTRATE GROWTH1* (*PROG1*) in chromosome 7 (30 cM), which is known to be a key gene for tillering in rice (Jin et al., 2008; Tan et al., 2008; Chen et al., 2019). Similar overlaps between genome regions highlighted by the ensemble models and QTL regions were also observed for ASI (Figure S5). These observed results indicated that multiple features highlighted in the ensemble models correspond to genome regions associated with several well-known key genes, implicating that effects associated with these genes contributed to the prediction performance of the ensembles.

## Discussion

### 1. Weighted ensemble performance is affected by the precision and diversity at the individual model level

This study investigated the possibility of improving the prediction performance using weighted ensemble approaches over the naïve ensemble-average methodology through weight optimisation. Overall, the weighted ensemble models did not always considerably improve the prediction performance for all the prediction scenarios considered in this study. They showed minor performance improvement in several scenarios compared to the naïve ensemble model (Figure 2, S2; Table S2). However, cumulative effects of modest improvements in predictive accuracy over multiple cycles of a breeding program can have substantial cumulative effects, as reported in Messina et al. (2023). Hence, the modest improvements demonstrated by the weighted ensemble models can contribute to the long-term acceleration of genetic gain.

In a number of cases, the three weighted ensemble-average models frequently achieved higher prediction performance for DTA and TILN than the naïve ensemble-average model across the datasets (Figure 2, S2; Table S2). In contrast, the prediction accuracy and error results from the naïve ensemble-average model were also observed within the results of the weighted ensemble-average models, particularly for ASI. The observed result indicates that the proposed weighted ensemble-average models were more effective in finding improvements over the naïve ensemble-average model for DTA and TILN rather than ASI. The equally weighted scenario of the naïve ensemble model might have already been close to the optimal weights for ASI in both datasets. Therefore, the weighted ensemble models could not find improvements in the weights for the individual prediction models. We clarify that the naïve ensemble-average model is a specific weighting pattern in weighted ensemble-average models, where the contributing individual models are given equal weight.

The lower accuracy in the inferred standing variation of the trait genetic architecture at the individual model level might have reduced the predictive ability of the weighted ensemble-average models for ASI. Such lower accuracy might have been attributed to the potential complexity of the genetic architecture. ASI is a secondary trait controlled by two quantitative flowering time traits (DTA and DTS) (Lu et al., 2010; Mhike et al., 2012; Messina et al., 2019), potentially increasing the complexity level of nonlinearity in the regulatory network. The component of nonlinearity was emphasised by the increased weights of the machine learning models for ASI in both datasets (Figure 3; Table S3), since machine learning models primarily focus on capturing complex nonlinear interactions between SNP genomic markers. However, the complex genetic architecture of ASI might have challenged the accurate detection of key genomic marker regions and their interactions by the individual genomic prediction models represented as one possible view of the standing variation of the trait genetic architecture rather than a complete one, including the machine learning models in this study. Such complexity in the regulatory network can also be further strengthened by the high sensitivity of ASI to genotype-by-environment (GxE) interactions (Buckler et al., 2009; Silva et al., 2022). Detailed environmental information can help prediction models learn key predictive patterns in environmental factors, possibly mitigating the negative effects of the complexity. However, the datasets used in this study did not include details of the environmental attributes. The difficulty in segregating environmental factors from the genetic architecture might have been amplified. The precision of the inferred standing variation of the trait genetic architecture at the individual model level and thereby their prediction performance might have been reduced (Cooper et al., 2022; Arenas et al., 2025). Since the prediction information from the individual level might have been inaccurate, it might have been difficult for the weighted ensemble-average models to optimise and diversify weights (Figure 3; Table S3). Consequently, the proposed weighted ensemble-average models might have failed to improve the prediction performance (Figure 2, S2; Table 1, S2) (Holland, 2007; Yang et al., 2011). Hence, high accuracy in the captured standing variation of the trait genetic architecture at the individual model level can be a critical factor in improving the prediction performance of the weighted ensemble-average models. The inferred standing variation of the trait genetic architecture can be verified through comparative analysis using the true trait genetic architecture defined from simulation datasets as a future research area.

The diversity of the individual genomic prediction models can be another factor influencing the prediction performance of ensembles. The inferred standing variation of the genetic architecture of the target traits from the individual genomic prediction models for the TeoNAM dataset was more diverse compared to that for the MaizeNAM dataset (Figure 5, S5). As implied by the Diversity Prediction Theorem (Hong and Page, 2004; Page, 2007, 2014, 2018), higher information diversity reduces the ensemble error by offsetting the weaknesses of individual genomic prediction models (Washburn et al., 2025). Assuming that the captured standing variation of the genetic architecture of DTA was more accurate than ASI, as discussed above, this higher diversity of the individual genomic prediction models at the genomic level might have contributed to the greater improvement in prediction performance with the weighted ensembles for DTA in the TeoNAM dataset compared to the MaizeNAM dataset. The diversity analysis using the two flowering time and tillering traits from two NAM datasets revealed that the diverse standing variation of the trait genetic architecture, in combination with the accuracy of the individual genomic prediction models, can be critical factors influencing the potential to amplify the predictive ability of ensembles through weight optimisation. The novelty of our study arises from analysing the association between the performance improvement level of the weighted ensemble models and the prediction performance and diversity at the individual prediction model level. Our study revealed scenarios when the weighted ensemble approaches can effectively improve prediction performance using the framework of the Diversity Prediction Theorem for crop breeding applications, which was not reported in previous weighted ensemble model studies (Kick and Washburn, 2023; Meher et al., 2025).

### 2. Multiple optimum weight combinations imply the absence of a consistent winner among the weighted ensembles

The prediction performance of the proposed weighted ensemble approaches was similar across the traits and datasets (Figure 2, S3). Strong positive associations between the weighted ensemble approaches were observed at both predicted phenotype and genomic marker effect levels (Figure 4, S4). Despite such high similarity at both levels, the weights assigned to each individual genomic prediction model were diverse, especially for DTA (Figure 3; Table S3). This suggests that multiple weight combinations can optimise prediction performance under the prediction scenarios in this study. The observed prediction performance among the weighted ensemble-average models implies the No Free Lunch Theorem (Wolpert and Macready, 1997), stating that the average performance of each prediction model becomes equivalent to others across all prediction scenarios. The effect of the No Free Lunch Theorem for genomic prediction in crop breeding has been widely observed and discussed at the individual genomic prediction level (Heslot et al., 2012; Haile et al., 2021; Merrick et al., 2021; Lourenço et al., 2024; Montesinos-Lopez et al., 2024; Crossa et al., 2025). The results from this study emphasise the No Free Lunch Theorem for genomic prediction in crop breeding applications at a higher prediction performance level by demonstrating no “best” weighted ensemble-average model when using similar objective functions.

The integration of the standing variation of the trait genetic architecture into ensemble models as prior knowledge can be an alternative approach to mitigate the effect of the No Free Lunch Theorem. The customisation of the prediction model into a problem-specific structure can mitigate the effect of the theorem (Montgomery, 2002). Traits are regulated by complex interactions between genetic markers, forming a gene network (Hammer et al., 2006; Cooper et al., 2009). Hence, inferring the standing variation for the trait genetic architecture in the form of a gene network model and integrating such information as prior knowledge can effectively supervise genomic prediction models to be problem-specific, especially when the training set size is small (Von Rueden et al., 2021). The gene network can be represented as a graph consisting of nodes and edges, and graph-oriented deep learning models such as graph neural networks can directly integrate such graphical information into their prediction mechanisms to predict phenotypes (Cooper et al., 2005; Xing et al., 2022). The incorporation of graph-based models into weighted ensemble-average models can be investigated to identify opportunities for further enhancing prediction performance.

### 3. A combined pipeline of hyperparameter tuning and weight optimisation may improve prediction performance

The diversity of individual genomic prediction models has been demonstrated and discussed as a critical factor in improving the prediction performance of the proposed weighted ensemble-average models in this study (Figures 3, 4, S4). However, the current weighted ensemble approaches do not explicitly coordinate between the individual genomic prediction models and the weighting algorithms to maximise the diversity without compromising the prediction accuracy, hindering the potential to reach the global optimum. As one approach for addressing this problem, the hyperparameters of the developed individual genomic prediction models can be tuned to maximise the improvement level of prediction performance using weight optimisation algorithms. The simultaneous optimisation of both weights and hyperparameters has improved prediction performance in several research studies (Lévesque et al., 2016; Wenzel et al., 2020; Shahhosseini et al., 2022). Hence, the prediction mechanisms of each individual genomic prediction model can be further adjusted to increase the diversity through hyperparameter tuning, combined with the weight optimisation incorporated into the weighted ensemble-average model (Shahhosseini et al., 2022).

The combined prediction pipeline can be created by seamlessly connecting the individual genomic prediction models and the weighted ensemble-average models by updating Equation (2) as follows:

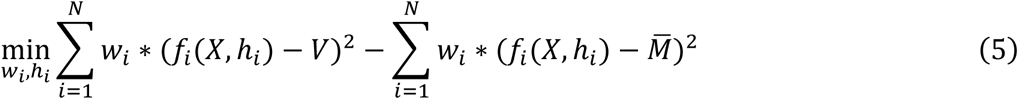

where *f_i_*(·) is an individual prediction model, *X* is the vector of genomic marker values and *h_i_* is the set of hyperparameters related to the prediction model (*f_i_*(·)). The hyperparameters of each individual genomic prediction model can be optimised using the validation set as well as the weights allocated to each. This pipeline can simultaneously optimise hyperparameters to diversify genomic prediction models with the consideration of the subsequent weight adjustment process, potentially further improving prediction performance. This is a promising area for further investigation that is highlighted by the current investigation of weighted ensemble prediction.

## Conclusion

In this study, the prediction performance of the three weighted ensemble-average models based on different optimisation strategies (linear transformation, Nelder-Mead and Bayesian algorithms) was evaluated against the prediction performance of the naïve ensemble-average model that calculates the arithmetic mean with equal weights. In agreement with the expectations of the Diversity Prediction Theorem, the naïve ensemble-average model performed well across traits and datasets. Our experimental results indicate the potential for performance improvement using the proposed weight optimisation approaches for several prediction scenarios.

However, the observed improvements were not achieved for all traits and dataset scenarios, suggesting opportunities for further investigation. We suggest that the weighted ensemble approaches can be further refined by developing a pipeline that can optimise both weights and hyperparameters of the individual genomic prediction models simultaneously.

## Data and code availability

The TeoNAM dataset (Chen et al., 2019) used in this study is located at https://datacommons.cyverse.org/browse/iplant/home/shared/panzea/genotypes/GBS/TeosinteNAM for the genotypes and https://doi.org/1-0.25386/genetics.9250682 for the phenotypes. The MaizeNAM dataset (Buckler et al., 2009) used in this study is located at https://cbsusrv04.tc.cornell.edu/users/panzea/download.aspx?filegroupid=10 for the genotypes and phenotypes. The computational tool, EasiGP, used in this study is located at https://github.com/ShunichiroT/EasiGP.

## Author contribution

ST designed the experiment, analysed data, visualised the results and wrote the manuscript. OP, MJW and JL provided feedback on the analysis, results and manuscript draft. MC formalised the concept, introduced the datasets, designed the experiment and provided feedback on the analysis, results and manuscript draft. All authors discussed the results and contributed to the writing and editing of the final version of the manuscript.

## Acknowledgments

We thank the National Computational Infrastructure (NCI) and the Research Computing Centre (RCC) at the University of Queensland for providing access to the High Performance Computing (HPC) machines.

## Supplementary Material

**Table S1:**
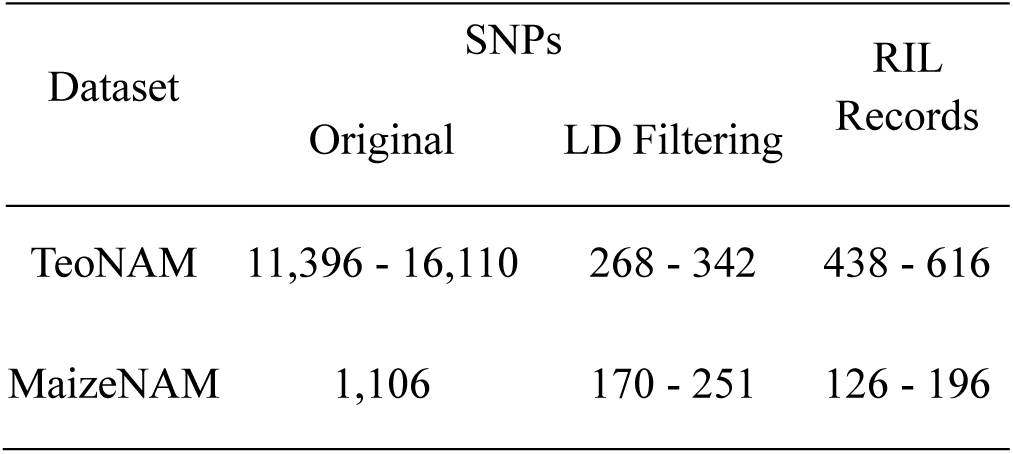
The total number of genomic markers (SNPs) at the original and after linkage disequilibrium (LD) filtering and the records of recombinant inbred lines (RILs) for the TeoNAM and MaizeNAM datasets.

| Dataset | SNPs |  | RIL<br>Records |
| --- | --- | --- | --- |
|  | Original | LD Filtering |  |
| TeoNAM | 11,396 - 16,110 | 268 - 342 | 438 - 616 |
| MaizeNAM | 1,106 | 170 - 251 | 126 - 196 |

**Table S2:**
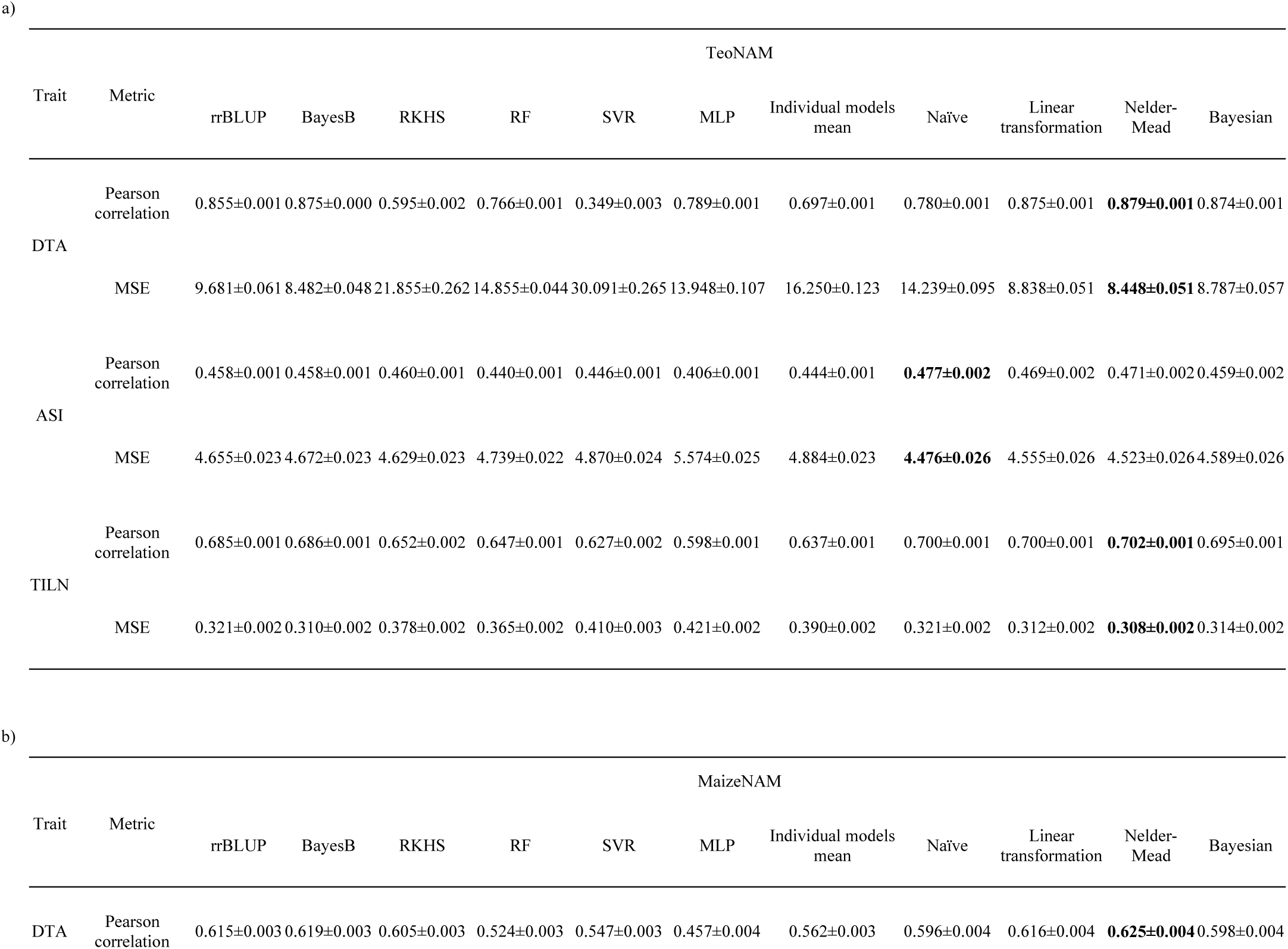
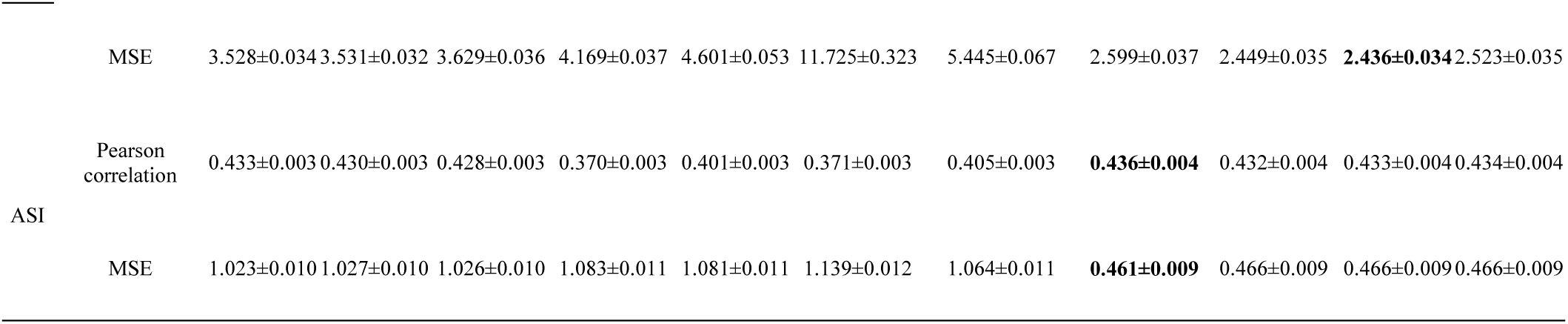
The median prediction accuracy (Pearson correlation) and error (mean squared error; MSE) of each ensemble model in the days to anthesis (DTA), anthesis and silking interval (ASI) and tiller number per plant (TILN) traits in the a) TeoNAM and b) MaizeNAM datasets. The TILN trait was recorded only in the TeoNAM dataset. The prediction performance was measured across 2,500 prediction scenarios for the TeoNAM dataset and 1,250 prediction scenarios for the MaizeNAM dataset, respectively, for each trait. The sign “±” indicates the standard error of the corresponding mean weight value. For prediction accuracy, the highest value in each prediction scenario was highlighted in bold, while for prediction error, the lowest value in each prediction scenario was highlighted in bold. This table corresponds to the violin plots in Figure 2 and S2.

**Table S3:** The mean weight and standard error values allocated to the six individual genomic prediction models (rrBLUP, BayesB, RKHS, RF, SVR and MLP) by the three optimisation approaches (linear transformation, Nelder-Mead and Bayesian ensemble-average approaches), along with naïve ensemble-average approach as the reference, for a) the days to anthesis (DTA), b) anthesis and silking interval (ASI) and c) tiller number per plant (TILN) traits in the TeoNAM and MaizeNAM datasets. The TILN trait was recorded only in the TeoNAM dataset. The weights were extracted from 2,500 prediction scenarios for the TeoNAM dataset and 1,250 prediction scenarios for the MaizeNAM dataset for each trait. The sign “±” indicates the standard error of the corresponding mean weight value. This table corresponds to Figure 3.

|  |  |  |  |  |  |  |  |  |
| --- | --- | --- | --- | --- | --- | --- | --- | --- |
| a) | DTA |  |  |  |  |  |  |  |
|  | TeoNAM |  |  |  | MaizeNAM |  |  |  |
|  | Naïve | Linear transformation | Nelder-Mead | Bayesian | Naïve | Linear transformation | Nelder-Mead | Bayesian |
| rrBLUP | 0.17±0 | 0.29±0.0017 | 0.19±0.0027 | 0.12±0.0039 | 0.17±0 | 0.22±0.0007 | 0.24±0.0038 | 0.03±0.0075 |
| BayesB | 0.17±0 | 0.30±0.0040 | 0.63±0.0028 | 0.37±0.0049 | 0.17±0 | 0.22±0.0057 | 0.31±0.0038 | 0.25±0.0079 |
| RKHS | 0.17±0 | 0.08±0.0029 | 0.03±0.0013 | 0.32±0.0009 | 0.17±0 | 0.18±0.0032 | 0.13±0.0034 | 0.47±0.0057 |
| RF | 0.17±0 | 0.23±0.0023 | 0.13±0.0024 | 0.15±0.0025 | 0.17±0 | 0.16±0.0010 | 0.15±0.0035 | 0.04±0.0057 |
| SVR | 0.17±0 | 0.02±0.0004 | 0.01±0.0003 | 0.01±0.0004 | 0.17±0 | 0.13±0.0009 | 0.08±0.0029 | 0.04±0.0043 |
| MLP | 0.17±0 | 0.07±0.0008 | 0.02±0.0011 | 0.04±0.0005 | 0.17±0 | 0.09±0.0029 | 0.08±0.0020 | 0.16±0.0026 |
| b) | ASI |  |  |  |  |  |  |  |
|  | TeoNAM |  |  |  | MaizeNAM |  |  |  |
|  | Naïve | Linear transformation | Nelder-Mead | Bayesian | Naïve | Linear transformation | Nelder-Mead | Bayesian |
| rrBLUP | 0.17±0 | 0.16±0.0043 | 0.17±0.0024 | 0.13±0.0050 | 0.17±0 | 0.18±0.0063 | 0.25±0.0034 | 0.12±0.0095 |
| BayesB | 0.17±0 | 0.16±0.0052 | 0.15±0.0025 | 0.20±0.0046 | 0.17±0 | 0.17±0.0096 | 0.19±0.0033 | 0.36±0.0087 |
| RKHS | 0.17±0 | 0.16±0.0040 | 0.10±0.0025 | 0.10±0.0040 | 0.17±0 | 0.17±0.0039 | 0.09±0.0033 | 0.06±0.0063 |
| RF | 0.17±0 | 0.20±0.0047 | 0.20±0.0028 | 0.19±0.0046 | 0.17±0 | 0.16±0.0069 | 0.17±0.0034 | 0.14±0.0078 |
| SVR | 0.17±0 | 0.13±0.0057 | 0.28±0.0019 | 0.28±0.0056 | 0.17±0 | 0.16±0.0070 | 0.13±0.0033 | 0.16±0.0068 |
| MLP | 0.17±0 | 0.18±0.0021 | 0.10±0.0026 | 0.10±0.0021 | 0.17±0 | 0.16±0.0066 | 0.17±0.0035 | 0.16±0.0066 |
| c) | TILN |  |  |  |  |  |  |  |
|  | TeoNAM |  |  |  | MaizeNAM |  |  |  |
|  | Naïve | Linear transformation | Nelder-Mead | Bayesian | Naïve | Linear transformation | Nelder-Mead | Bayesian |
| rrBLUP | 0.17±0 | 0.18±0.0035 | 0.17±0.0024 | 0.10±0.0047 | - | - | - | - |
| BayesB | 0.17±0 | 0.22±0.0051 | 0.35±0.0025 | 0.43±0.0056 | - | - | - | - |
| RKHS | 0.17±0 | 0.13±0.0022 | 0.07±0.0022 | 0.05±0.0032 | - | - | - | - |
| RF | 0.17±0 | 0.17±0.0033 | 0.15±0.0024 | 0.14±0.0033 | - | - | - | - |
| SVR | 0.17±0 | 0.13±0.0038 | 0.13±0.0021 | 0.15±0.0037 | - | - | - | - |
| MLP | 0.17±0 | 0.18±0.0024 | 0.13±0.0023 | 0.13±0.0024 | - | - | - | - |

**Figure S1:**
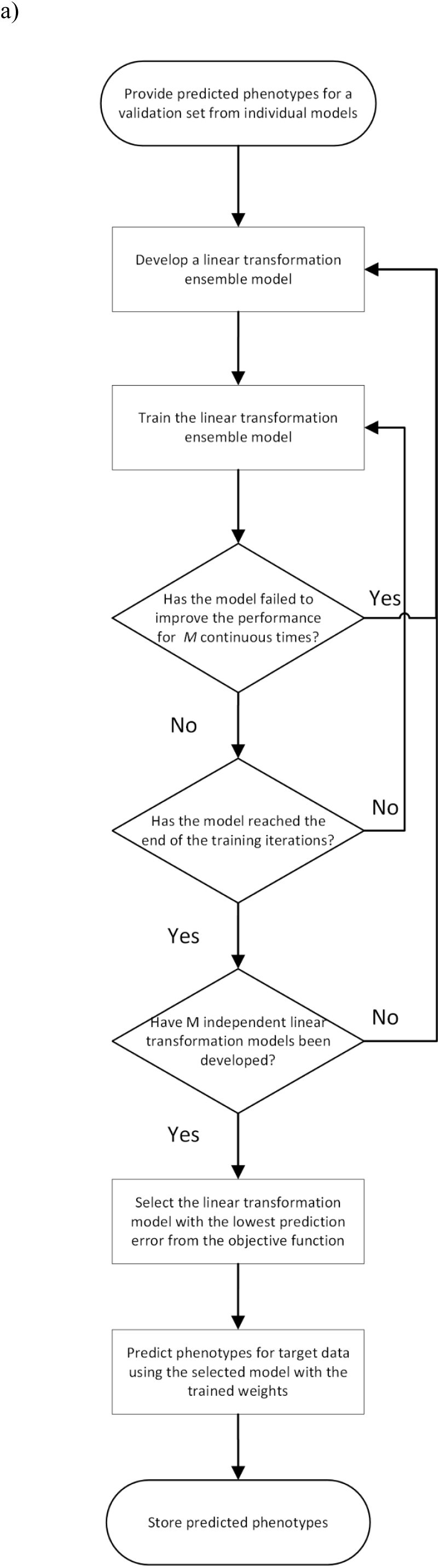
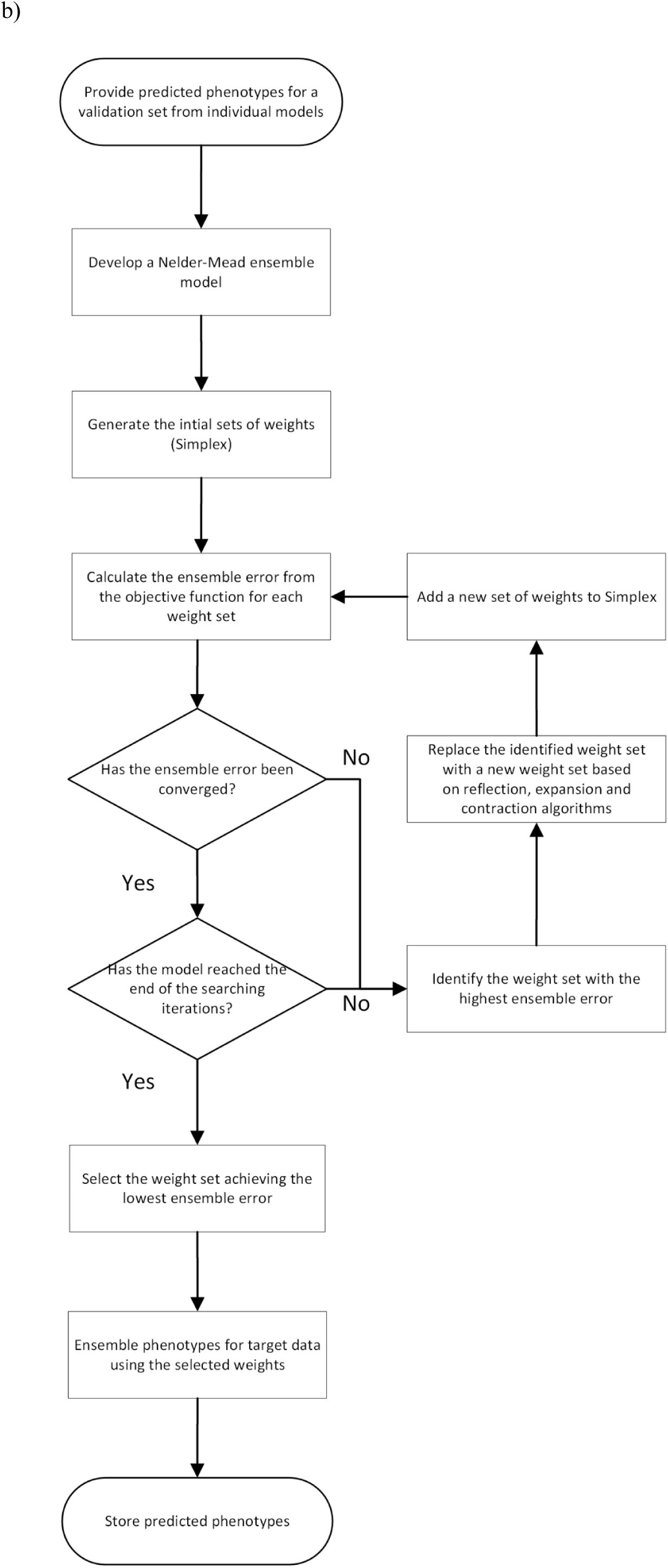
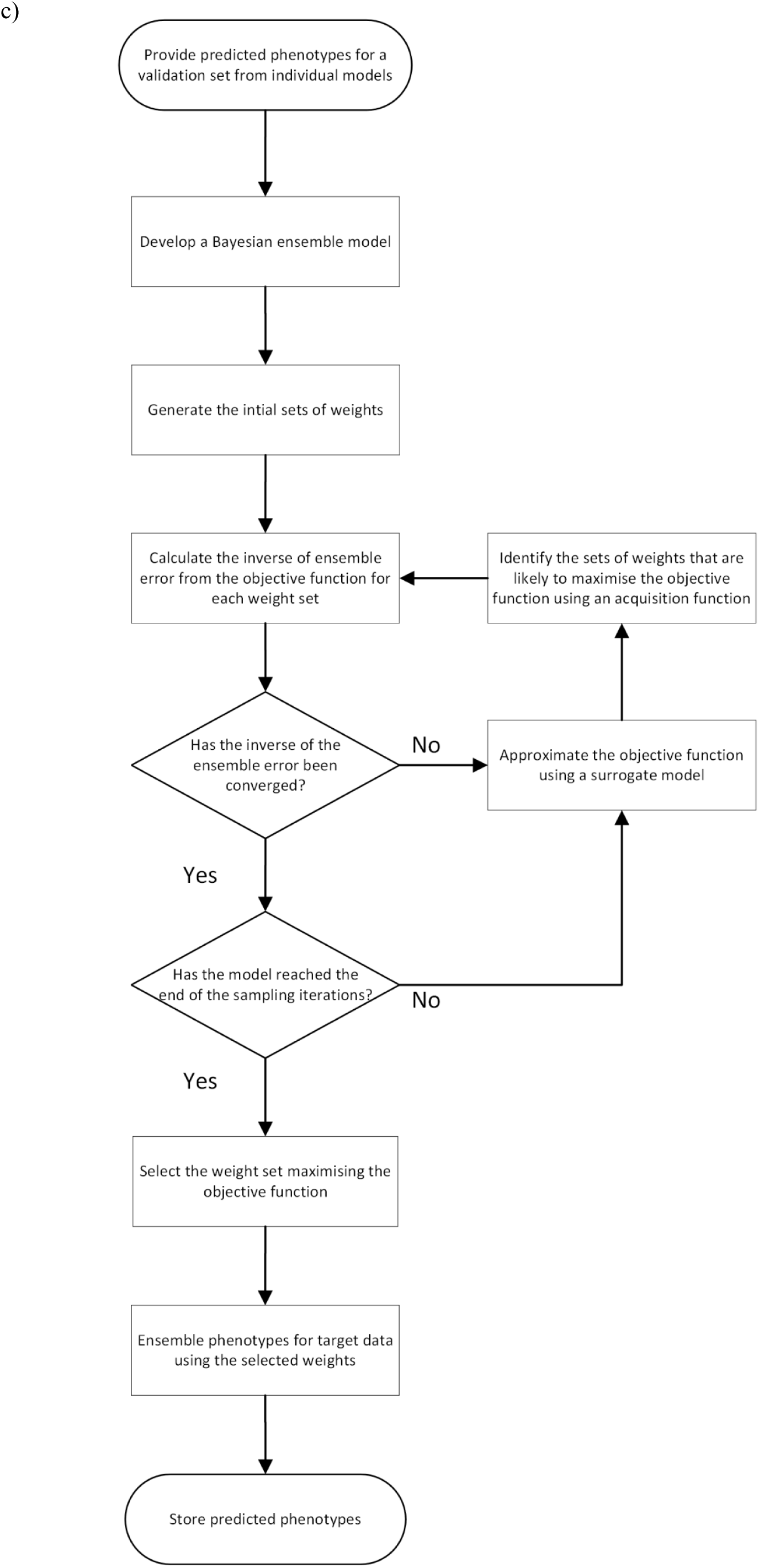
Flowchart for the weighted ensemble models used in this study; the a) linear transformation ensemble, b) Nelder-Mead ensemble and c) Bayesian ensemble models.

**Figure S2:**
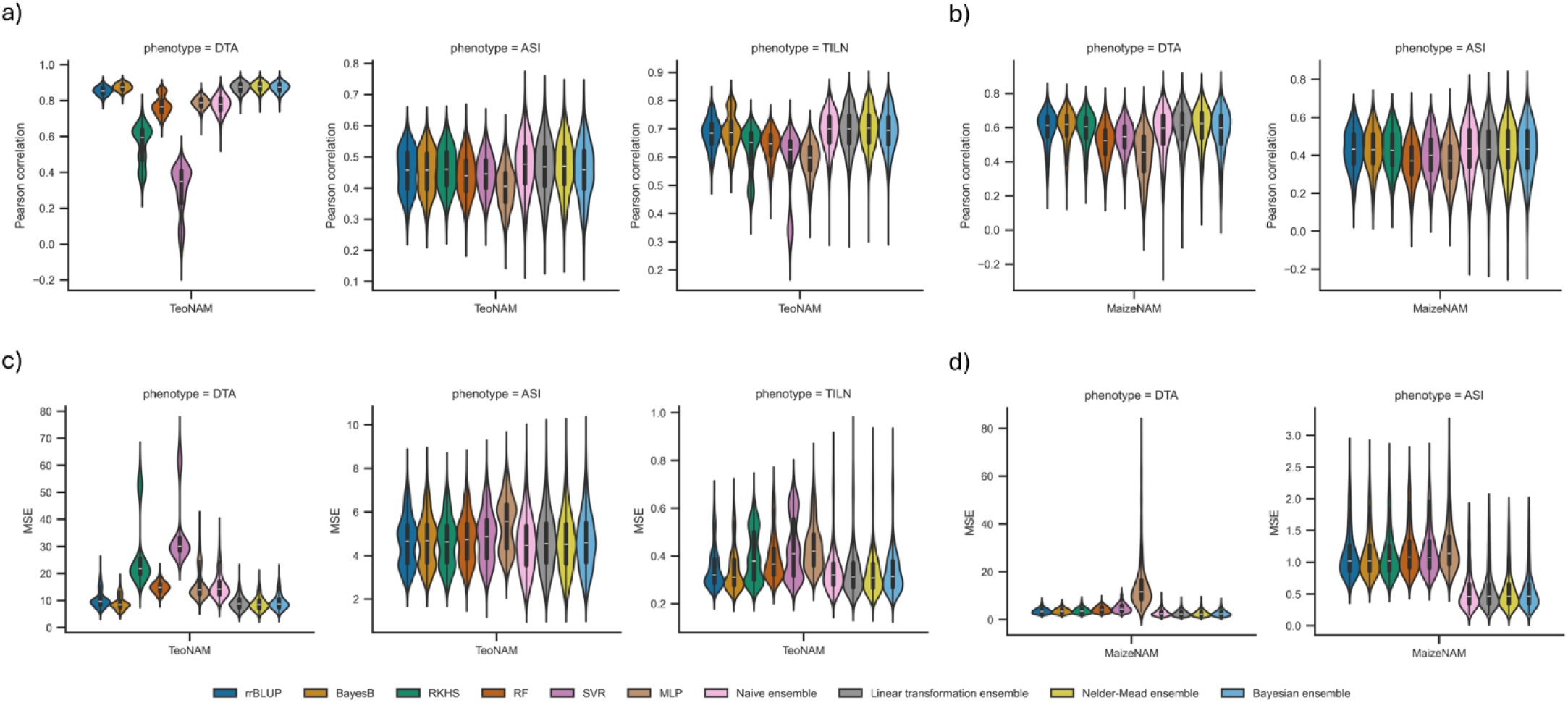
Comparison of median prediction performance between the individual genomic prediction models (rrBLUP, BayesB, RKHS, RF, SVR and MLP) and the ensemble-average models (naïve ensemble, linear transformation ensemble, Nelder-Mead ensemble and Bayesian ensemble) for the days to anthesis (DTA), anthesis and silking interval (ASI) and tiller number per plant (TILN) traits in the TeoNAM and MeizeNAM dataset. The TILN trait was recorded only in the TeoNAM dataset. Pearson correlation and mean squared error (MSE) were used as performance metrics; a) Pearson correlation for the TeoNAM dataset, b) Pearson correlation for the MaizeNAM dataset, c) MSE for the TeoNAM dataset and d) MSE for the MaizeNAM dataset. The prediction metrics were calculated from 2,500 prediction scenarios for the TeoNAM dataset and 1,250 prediction scenarios for the MaizeNAM dataset for each trait. The width of the violins represents the distribution of performance metrics. The white horizontal lines on the black box plots show the median value for each metric. The whiskers extend 1.5 times the interquartile range. The median value of each metric is recorded in Table S2.

**Figure S3:**
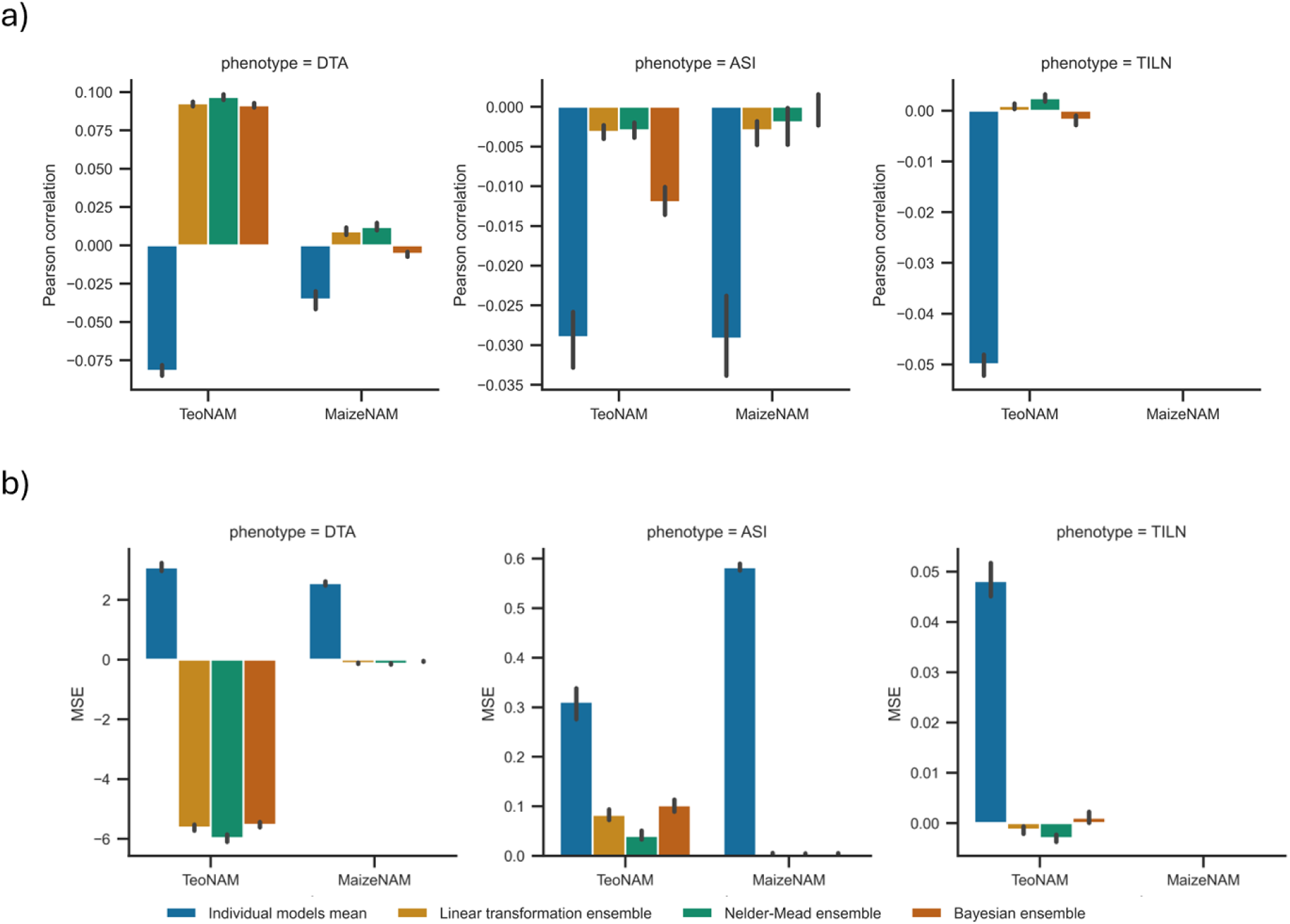
Comparison of median prediction performance difference with the naïve ensemble-average model between the mean metric value of the individual genomic prediction models (rrBLUP, BayesB, RKHS, RF, SVR and MLP) and the weighted ensemble-average models (linear transformation ensemble, Nelder-Mead ensemble and Bayesian ensemble). The performance of the prediction models was compared for the days to anthesis (DTA), anthesis and silking interval (ASI) and tiller number per plant (TILN) in the TeoNAM and MaizeNAM datasets. The TILN trait was recorded only in the TeoNAM dataset. The prediction performance was measured using the two metrics: a) Pearson correlation and b) mean squared error (MSE). For Pearson correlation, positive values indicate that corresponding prediction models achieved higher prediction accuracy than the naïve ensemble average model and vice versa. For MSE, negative values indicate that the corresponding prediction models achieved lower prediction error than the naïve ensemble average model and vice versa. The prediction metrics were calculated from 2,500 prediction scenarios for the TeoNAM dataset and 1,250 prediction scenarios for the MaizeNAM dataset for each trait. The black vertical line on each bar shows the error bar of the confidence interval. This figure corresponds to Figure 2.

**Figure S4:**
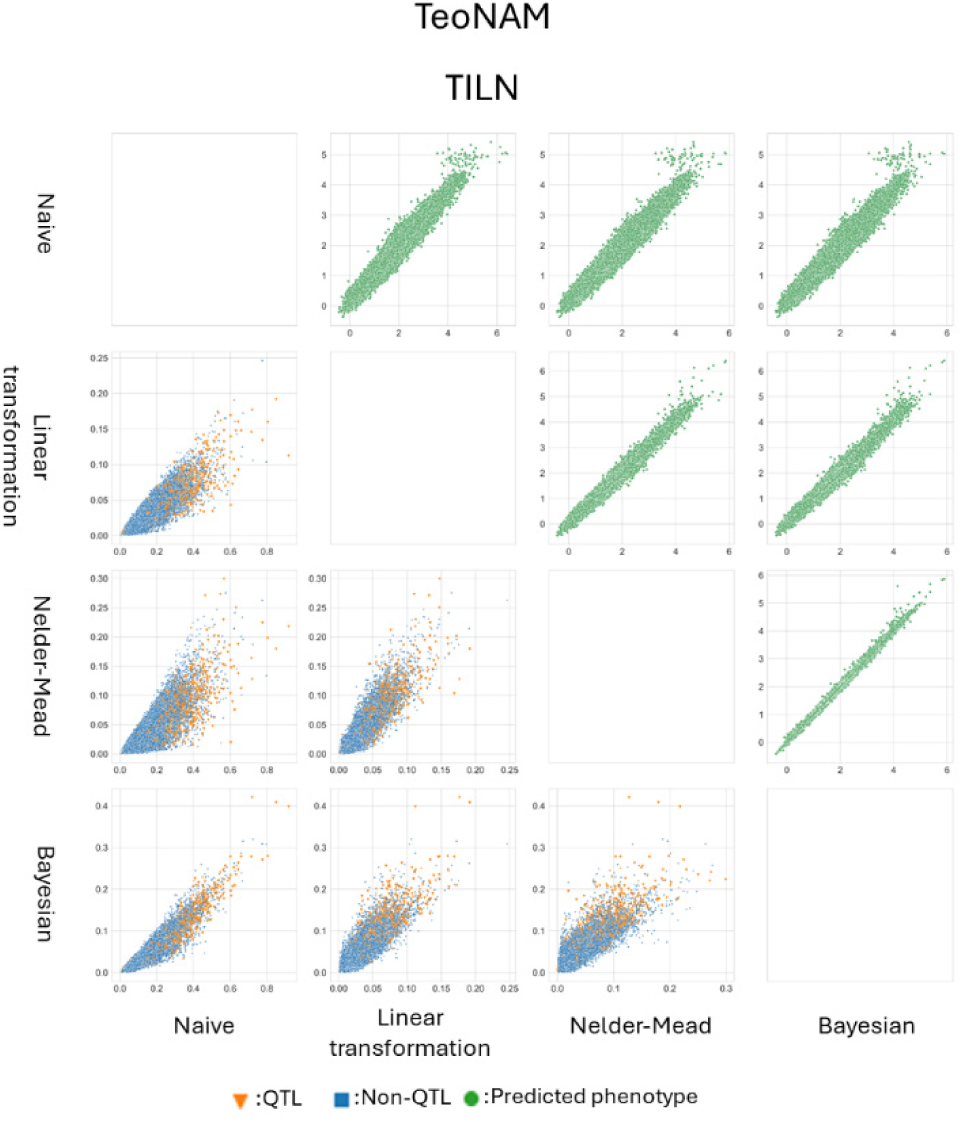
Pairwise comparisons of the ensemble models (naïve ensemble, linear transformation, Nelder-Mead and Bayesian) for the tiller number per plant (TILN) trait across all the prediction scenarios for the TeoNAM (2,500 prediction scenarios). Within each scatter plot matrix, the ensemble models were compared at the predicted phenotypes (top right triangle) and genomic marker effects (the bottom left triangle) levels. The green dots represent a pair of predicted phenotypes for RIL samples in the test sets for each prediction scenario. The blue squares and orange triangles represent pairs of inferred effects of genomic markers in each sample scenario for non-QTL and QTL, respectively. Non-QTL and QTL markers were identified by Chen et al. (2019).

**Figure S5:**
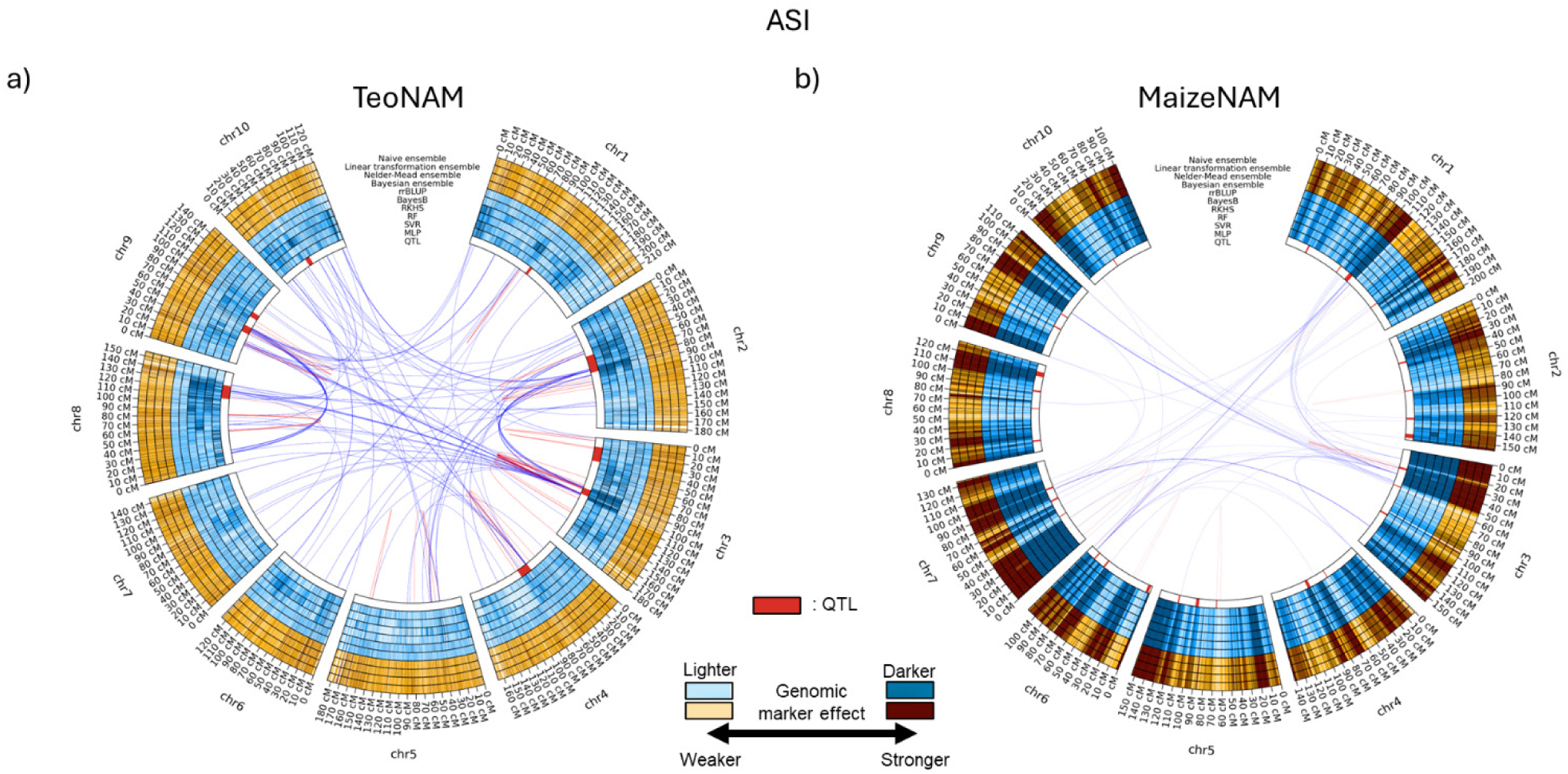
Circos plots for the anthesis and silking interval (ASI) in the a) TeoNAM and b) MaizeNAM datasets. The innermost (QTL) ring shows the QTL gene regions estimated by Chen et al. (2019) for the TeoNAM dataset and Buckler et al. (2009) for the MaizeNAM dataset. The blue rings indicate genomic marker effects across the gene regions estimated by MLP, SVR, RF, RKHS, BayesB and rrBLUP. The subsequent four orange rings outwards are the genomic marker effects estimated by Bayesian, Nelder-Mead, linear transformation and naïve ensemble. The darkness level of the blue and orange colours indicates the strength of the genomic marker effects, sectioned into ten levels using the quantiles. Darker colours represent higher genomic marker effect sizes. The red and blue lines between genome regions show the genomic marker interaction effects calculated from pairwise Shapley scores from RF (top 0.01%; red = within chromosome and blue = between chromosomes).

## References

Arenas S, Djabali Y, Rincent R, Cubry P, Martin ML, Blein-Nicolas M, Laplaze L, Schneider H, Grondin A. 2025. Modeling plant phenotypic plasticity and its underlying genetic architecture: a comparative study. Journal of Experimental Botany. p. eraf013. 10.1093/jxb/eraf013.

Bernardo R, Yu J. 2007. Prospects for genomewide selection for quantitative traits in maize. Crop Science 47:1082–1090. 10.2135/cropsci2006.11.0690.

Bellman R. 1957. Dynamic programming. Princeton University Press Princeton. pp. 24–73.

Breiman L. 2001. Random forests. Machine Learning. 45:5–32. 10.1023/A:1010933404324.

Buckler ES, Holland JB, Bradbury PJ, Acharya CB, Brown PJ, Browne C, Ersoz E, Flint-Garcia S, Garcia A, Glaubitz JC et al. 2009. The genetic architecture of maize flowering time. Science. 325:714–718. 10.1126/science.1174276.

Chang CC, Chow CC, Tellier LC, Vattikuti S, Purcell SM, Lee JJ. 2015. Second-generation plink: rising to the challenge of larger and richer datasets. GigaScience. 4. 10.1186/s13742015-0047-8.

Chen Q, Yang CJ, York AM, Xue W, Daskalska LL, DeValk CA, Krueger KW, Lawton SB, Spiegelberg BG, Schnell JM et al. 2019. Teonam: A nested association mapping population for domestication and agronomic trait analysis in maize. Genetics. 213:1065–1078. 10.1534/genetics.119.302594.

Cooper M, Messina CD, Tang T, Gho C, Powell OM, Podlich DW, Technow F, Hammer GL. 2022. Predicting genotype× environment× management (g× e× m) interactions for the design of crop improvement strategies: integrating breeder, agronomist, and farmer perspectives. Plant breeding reviews. 46:467–585. 10.1002/9781119874157.ch8.

Cooper M, Podlich DW, Smith OS. 2005. Gene-to-phenotype models and complex trait genetics. Australian Journal of Agricultural Research. 56:895–918. 10.1071/AR05154

Cooper M, Tomura S, Wilkinson MJ, Powell O, Messina CD. 2025. Breeding perspectives on tackling trait genome-to-phenome (g2p) dimensionality using ensemble-based genomic prediction. Theoretical and Applied Genetics. 138:172. 10.1007/s00122-025-04960-6.

Cooper M, van Eeuwijk FA, Hammer GL, Podlich DW, Messina C. 2009. Modeling qtl for complex traits: detection and context for plant breeding. Current opinion in plant biology. 12:231–240. 10.1016/j.pbi.2009.01.006.

Cooper M, Voss-Fels KP, Messina CD, Tang T, Hammer GL. 2021. Tackling g× e× m interactions to close on-farm yield-gaps: creating novel pathways for crop improvement by predicting contributions of genetics and management to crop productivity. Theoretical and Applied Genetics. 134:1625–1644. 10.1007/s00122-021-03812-3.

Crossa J, Montesinos-Lopez OA, Costa-Neto G, Vitale P, Martini JW, Runcie D, Fritsche-Neto R, Montesinos-Lopez A, Pérez-Rodríguez P, Gerard G et al. 2024. Machine learning algorithms translate big data into predictive breeding accuracy. Trends in Plant Science. 10.1016/j.tplants.2024.09.011.

Crossa J, Pérez-Rodríguez P, Cuevas J, Montesinos-López O, Jarquín D, de los Campos G, Burguengo J, González-Camacho JM, Pérez-Elizalde S, Beyene Y et al. 2017. Genomic selection in plant breeding: methods, models, and perspectives. Trends in plant science. 22:961–975. 10.1016/j.tplants.2017.08.011.

de los Campos G, Gianola D, Rosa GJ. 2009. Reproducing kernel hilbert spaces regression: a general framework for genetic evaluation. Journal of animal science. 87:1883–1887. 10.2527/jas.2008-1259.

Doebley J, Stec A, Hubbard L. 1997. The evolution of apical dominance in maize. Nature. 386:485–488. 10.1038/386485a0.

Dong Z, Danilevskaya O, Abadie T, Messina C, Coles N, Cooper M. 2012. A gene regulatory network model for floral transition of the shoot apex in maize and its dynamic modeling. Plos One. 10.1371/journal.pone.0043450.

Drucker H, Burges CJ, Kaufman L, Smola A, Vapnik V. 1996. Support vector regression machines. Advances in neural information processing systems. 9.

Endelman JB. 2011. Ridge regression and other kernels for genomic selection with r package rrblup. The Plant Genome. 4:250–255. 10.3835/plantgenome2011.08.0024.

Escamilla DM, Li D, Negus KL, Kappelmann KL, Kusmec A, Vanous AE, Schnable PS, Li X, Yu J. 2025. Genomic selection: Essence, applications, and prospects. The Plant Genome. 18:e70053. 10.1002/tpg2.70053.

Gianola D, van Kaam JBCHM. 2008. Reproducing kernel hilbert spaces regression methods for genomic assisted prediction of quantitative traits. Genetics. 178:2289–2303. 10.1534/genetics.107.084285.

Haile TA, Walkowiak S, N’Diaye A, Clarke JM, Hucl PJ, Cuthbert RD, Knox RE, Pozniak CJ. 2021. Genomic prediction of agronomic traits in wheat using different models and cross-validation designs. Theoretical and Applied Genetics. 134:381–398. 10.1007/s00122-020-03703-z.

Hammer G, Cooper M, Tardieu F, Welch S, Walsh B, van Eeuwijk F, Chapman S, Podlich D. 2006. Models for navigating biological complexity in breeding improved crop plants. Trends in Plant Science 11(12):587–593. 10.1016/j.tplants.2006.10.006.

Hammer GL, McLean G, Kholová J, van Oosterom E. 2023. Modelling the dynamics and phenotypic consequences of tiller outgrowth and cessation in sorghum. in silico Plants. 5:diad019. 10.1093/insilicoplants/diad019.

Heffner EL, Sorrells ME, Jannink JL. 2009. Genomic selection for crop improvement. Crop Science. 49:1–12. 10.2135/cropsci2008.08.0512.

Heslot N, Yang HP, Sorrells ME, Jannink JL. 2012. Genomic selection in plant breeding: a comparison of models. Crop science. 52:146–160. 10.2135/cropsci2011.06.0297.

Holland JB. 2007. Genetic architecture of complex traits in plants. Current opinion in plant biology. 10:156–161. 10.1016/j.pbi.2007.01.003.

Hong L, Page SE. 2004. Groups of diverse problem solvers can outperform groups of highability problem solvers. Proceedings of the National Academy of Sciences. 101:16385–16389. 10.1073/pnas.0403723101.

Huang C, Sun H, Xu D, Chen Q, Liang Y, Wang X, Xu G, Tian J, Wang C, Li D et al. 2018. Zmcct9 enhances maize adaptation to higher latitudes. Proceedings of the National Academy of Sciences. 115:E334–E341. 10.1073/pnas.1718058115.

Hufford MB, Xu X, Van Heerwaarden J, Pyhäjärvi T, Chia JM, Cartwright RA, Elshire RJ, Glaubitz JC, Guill KE, Kaeppler SM et al. 2012. Comparative population genomics of maize domestication and improvement. Nature genetics. 44:808–811. 10.1038/ng.2309.

Ishwaran H. 2015. The effect of splitting on random forests. Machine learning. 99:75–118. 10.1007/s10994-014-5451-2.

Jin J, Huang W, Gao JP, Yang J, Shi M, Zhu MZ, Luo D, Lin HX. 2008. Genetic control of rice plant architecture under domestication. Nature genetics. 40:1365–1369. 10.1038/ng.247.

Kick DR, Washburn JD. 2023. Ensemble of best linear unbiased predictor, machine learning and deep learning models predict maize yield better than each model alone. in silico Plants. 5:diad015. 10.1093/insilicoplants/diad015.

Kingma DP, Ba J. 2014. Adam: A method for stochastic optimization. arXiv preprint arXiv:1412.6980. 10.48550/arXiv.1412.6980.

Krzywinski M, Schein J, Birol I, Connors J, Gascoyne R, Horsman D, Jones SJ, Marra MA. 2009. Circos: an information aesthetic for comparative genomics. Genome research. 19:1639–1645. 10.1126/science.1174276.

Lévesque JC, Gagné C, Sabourin R. 2016. Bayesian hyperparameter optimization for ensemble learning. arXiv preprint arXiv:1605.06394. 10.48550/arXiv.1605.06394.

Liang M, Cao S, Deng T, Du L, Li K, An B, Du Y, Xu L, Zhang L, Gao X et al. 2023. Mak: a machine learning framework improved genomic prediction via multi-target ensemble regressor chains and automatic selection of assistant traits. Briefings in Bioinformatics. 24:bbad043. 10.1093/bib/bbad043.

Liang M, Chang T, An B, Duan X, Du L, Wang X, Miao J, Xu L, Gao X, Zhang L et al. 2021a. A stacking ensemble learning framework for genomic prediction. Frontiers in genetics. 12:600040. 10.3389/fgene.2021.600040.

Liang M, Miao J, Wang X, Chang T, An B, Duan X, Xu L, Gao X, Zhang L, Li J et al. 2021b. Application of ensemble learning to genomic selection in chinese simmental beef cattle. Journal of Animal Breeding and Genetics. 138:291–299. 10.1111/jbg.12514.

Loshchilov I, Hutter F. 2017. Decoupled weight decay regularization. arXiv preprint arXiv:1711.05101. 10.48550/arXiv.1711.05101.

Lourenço VM, Ogutu JO, Rodrigues RA, Posekany A, Piepho HP. 2024. Genomic prediction using machine learning: a comparison of the performance of regularized regression, ensemble, instance-based and deep learning methods on synthetic and empirical data. BMC genomics. 25:152. 10.1186/s12864-023-09933-x.

Lu Y, Zhang S, Shah T, Xie C, Hao Z, Li X, Farkhari M, Ribaut JM, Cao M, Rong T et al. 2010. Joint linkage–linkage disequilibrium mapping is a powerful approach to detecting quantitative trait loci underlying drought tolerance in maize. Proceedings of the National Academy of Sciences. 107:19585– 19590. 10.1073/pnas.1006105107.

Lundberg SM, Lee SI. 2017. A unified approach to interpreting model predictions. Advances in neural information processing systems. 30.

McCormick RF, Truong SK, Rotundo J, Gaspar AP, Kyle D, Van Eeuwijk F, Messina CD. 2021. Intercontinental prediction of soybean phenology via hybrid ensemble of knowledge-based and datadriven models. in silico Plants. 3:diab004. 10.1093/insilicoplants/diab004.

McMullen MD, Kresovich S, Villeda HS, Bradbury P, Li H, Sun Q, Flint-Garcia S, Thornsberry J, Acharya C, Bottoms C et al. 2009. Genetic properties of the maize nested association mapping population. Science. 325:737–740. 10.1126/science.1174320.

Meher PK, Pradhan UK, Ray M, Gupta A, Parsad R, Gupta PK. 2025. Ensemble of bayesian alphabets via constraint weight optimization strategy improves genomic prediction accuracy. G3: Genes, Genomes, Genetics. 15:jkaf150. 10.1093/g3journal/jkaf150.

Merrick LF, Carter AH. 2021. Comparison of genomic selection models for exploring predictive ability of complex traits in breeding programs. The Plant Genome. 14:e20158. 10.1002/tpg2.20158.

Messina C, Garcia-Abadillo J, Powell O, Tomura S, Zare A, Ganapathysubramanian B, Cooper M. 2025. Toward a general framework for ai-enabled prediction in crop improvement. Theoretical and Applied Genetics. 138:151. 10.1007/s00122-025-04928-6.

Messina CD, Gho C, Hammer GL, Tang T, Cooper M. 2023. Two decades of harnessing standing genetic variation for physiological traits to improve drought tolerance in maize. Journal of experimental botany. 74:4847–4861. 10.1093/jxb/erad231.

Messina CD, Hammer GL, McLean G, Cooper M, van Oosterom EJ, Tardieu F, Chapman SC, Doherty A, Gho C. 2019. On the dynamic determinants of reproductive failure under drought in maize. in silico Plants. 1:diz003. 10.1093/insilicoplants/diz003.

Messina CD, Technow F, Tang T, Totir R, Gho C, Cooper M. 2018. Leveraging biological insight and environmental variation to improve phenotypic prediction: Integrating crop growth models (cgm) with whole genome prediction (wpg). European Journal of Agronomy 100:151–162. 10.1016/j.eja.2018.01.007.

Meuwissen TH, Hayes BJ, Goddard M. 2001. Prediction of total genetic value using genome-wide dense marker maps. genetics. 157:1819–1829. 10.1093/genetics/157.4.1819.

Mhike X, Okori P, Magorokosho C, Ndlela T. 2012. Validation of the use of secondary traits and selection indices for drought tolerance in tropical maize (zea mays l.). Afr. J. Plant Sci. 6:96–102. 10.5897/AJPS11.179.

Montesinos-Lopez A, Crespo-Herrera L, Dreisigacker S, Gerard G, Vitale P, Saint Pierre C, Govindan V, Tarekegn ZT, Flores MC, Pérez-Rodríguez P et al. 2024. Deep learning methods improve genomic prediction of wheat breeding. Frontiers in Plant Science. 15:1324090. 10.3389/fpls.2024.1324090.

Montgomery J. 2002. The no free lunch theorems for optimisation: An overview. The No Free Lunch Theorems, Evolution and Evolutionary Algorithms.

Nair V, Hinton GE. 2010. Rectified linear units improve restricted boltzmann machines. pp. 807–814.

Negus, KL, Li X, Welch SM, Yu J. 2024. The role of artificial intelligence in crop improvement. Advances in Agronomy 184:1–66. 10.1016/bs.agron.2023.11.001.

Nelder JA, Mead R. 1965. A simplex method for function minimization. The computer journal. 7:308– 313. 10.1093/comjnl/7.4.308.

Nogueira F. 2014. Bayesian Optimization: Open source constrained global optimization tool for Python.

Page SE. 2007. Making the difference: Applying a logic of diversity. Academy of Management Perspectives. 21:6–20. 10.5465/amp.2007.27895335.

Page SE. 2014. Where diversity comes from and why it matters? European Journal of Social Psychology. 44:267–279. 10.1002/ejsp.2016.

Page SE. 2018. The model thinker: What you need to know to make data work for you. Hachette UK.

Pérez P, de los Campos G. 2014. Genome-wide regression and prediction with the bglr statistical package. Genetics. 198:483–495. 10.1534/genetics.114.164442.

Powell OM, Barbier F, Voss-Fels KP, Beveridge C, Cooper M. 2022. Investigations into the emergent properties of gene-to-phenotype networks across cycles of selection: a case study of shoot branching in plants. in silico Plants. 4:diac006. 10.1093/insilicoplants/diac006.

Ramstein GP, Jensen SE, Buckler ES. 2019. Breaking the curse of dimensionality to identify causal variants in breeding 4. Theoretical and Applied Genetics. 132:559–567. 10.1007/s00122018-3267-3.

Riedelsheimer C, Technow F, Melchinger AE. 2012. Comparison of whole-genome prediction models for traits with contrasting genetic architecture in a diversity panel of maize inbred lines. BMC genomics. 13:1–9. 10.1186/1471-2164-13-452.

Rosenblatt F. 1962. Principles of neurodynamics. Perceptrons and the theory of brain mechanisms. 10.1037/h0042519.

Shahhosseini M, Hu G, Pham H. 2022. Optimizing ensemble weights and hyperparameters of machine learning models for regression problems. Machine Learning with Applications. 7:100251. 10.1016/j.mlwa.2022.100251.

Shapley LS. 1953. Stochastic games. Proceedings of the national academy of sciences. 39:1095–1100. 10.1073/pnas.39.10.1095.

Silva PC, Sanchez AC, Opazo MA, Mardones LA, Acevedo EA. 2022. Grain yield, anthesis-silking interval, and phenotypic plasticity in response to changing environments: Evaluation in temperate maize hybrids. Field Crops Research. 285:108583. 10.1016/j.fcr.2022.108583.

Singer S, Nelder J. 2009. Nelder-mead algorithm. Scholarpedia. 4:2928. 10.4249/scholarpedia.2928.

Studer AJ, Wang H, Doebley JF. 2017. Selection during maize domestication targeted a gene network controlling plant and inflorescence architecture. Genetics. 207:755–765.10.1534/genetics.117.300071.

Tan L, Li X, Liu F, Sun X, Li C, Zhu Z, Fu Y, Cai H, Wang X, Xie D et al. 2008. Control of a key transition from prostrate to erect growth in rice domestication. Nature genetics. 40:1360–1364. 10.1038/ng.197.

Technow F, Messina CD, Totir LR, Cooper M (2015) Integrating crop growth models with whole genome prediction through approximate Bayesian computation. PloS One 10:e0130855. 10.1371/journal.pone.0130855.

Tieleman T. 2012. Lecture 6.5-rmsprop: Divide the gradient by a running average of its recent magnitude. COURSERA: Neural networks for machine learning. 4:26.

Tomura S, Powell O, Wilkinson MJ, Cooper M. 2026. Ensemble-based genomic prediction for maize flowering-time improves prediction accuracy and reveals novel insights into trait genetic variation. G3: Genes, Genomes, Genetics. 10.1093/g3journal/jkag090.

Tomura S, Wilkinson MJ, Cooper M, Powell O. 2025a. Improved genomic prediction performance with ensembles of diverse models. G3: Genes, Genomes, Genetics. p. jkaf048. 10.1093/g3journal/jkaf048.

Tomura S, Wilkinson MJ, Powell O, Cooper M. 2025b. Ensemble analysis with interpretable genomic prediction (easigp): Computational tool for interpreting ensembles of genomic prediction models. The Plant Genome. 18:e70138. 10.1002/tpg2.70138.

Von Rueden L, Mayer S, Beckh K, Georgiev B, Giesselbach S, Heese R, Kirsch B, Pfrommer J, Pick A, Ramamurthy R et al. 2021. Informed machine learning–a taxonomy and survey of integrating prior knowledge into learning systems. IEEE Transactions on Knowledge and Data Engineering. 35:614–633. 10.1109/TKDE.2021.3079836.

Voss-Fels KP, Cooper M, Hayes BJ. 2019. Accelerating crop genetic gains with genomic selection. Theoretical and Applied Genetics. 132:669–686. 10.1007/s00122-018-3270-8.

Wallach D, Martre P, Liu B, Asseng S, Ewert F, Thorburn PJ, van Ittersum M, Aggarwal PK, Ahmed M, Basso B et al. 2018. Multimodel ensembles improve predictions of crop–environment– management interactions. Global change biology. 24:5072–5083. 10.1111/gcb.14411.

Wang Z, Wang H, Yu T, Zhang W, Han J, Li F. 2023. A multiple kernel ensemble approach for genomic prediction. volume 12609. pp. 324–336. SPIE. 10.1117/12.2671691.

Washburn JD, Varela JI, Xavier A, Chen Q, Ertl D, Gage JL, Holland JB, Lima DC, Romay MC, Lopez-Cruz M et al. 2025. Global genotype by environment prediction competition reveals that diverse modeling strategies can deliver satisfactory maize yield estimates. Genetics. 229:iyae195. 10.1093/genetics/iyae195.

Wenzel F, Snoek J, Tran D, Jenatton R. 2020. Hyperparameter ensembles for robustness and uncertainty quantification. Advances in Neural Information Processing Systems. 33:6514–6527.

Wisser RJ, Fang Z, Holland JB, Teixeira JE, Dougherty J, Weldekidan T, de Leon N, Flint-Garcia S, Lauter N, Murray SC et al. 2019. The genomic basis for short-term evolution of environmental adaptation in maize. Genetics. 213:1479–1494. 10.1534/genetics.119.302780.

Wolpert DH, Macready WG. 1997. No free lunch theorems for optimization. IEEE transactions on evolutionary computation. 1:67–82. 10.1109/4235.585893.

Xing X, Yang F, Li H, Zhang J, Zhao Y, Gao M, Huang J, Yao J. 2022. Multi-level attention graph neural network based on co-expression gene modules for disease diagnosis and prognosis. Bioinformatics. 38:2178–2186. 10.1093/bioinformatics/btac088.

Yan J, Xu Y, Cheng Q, Jiang S, Wang Q, Xiao Y, Ma C, Yan J, Wang X. 2021. Lightgbm: accelerated genomically designed crop breeding through ensemble learning. Genome biology. 22:1–24. 10.1186/s13059-021-02492-y.

Yang L. 2011. Classifiers selection for ensemble learning based on accuracy and diversity. Procedia Engineering. 15:4266–4270. 10.1016/j.proeng.2011.08.800.

Yu T, Zhang W, Han J, Li F, Wang Z, Cao C. 2021. An ensemble learning approach for predicting phenotypes from genotypes. pp. 382–389. IEEE. 10.1109/IUCC-CIT-DSCISmartCNS55181.2021.00068.

Zhao Q, Weber AL, McMullen MD, Guill K, Doebley J. 2011. Mads-box genes of maize: frequent targets of selection during domestication. Genetics research. 93:65–75. 10.1017/S0016672310000509.

